# Decoding regulatory specificity of human ribosomal proteins

**DOI:** 10.1101/2021.03.27.437302

**Authors:** Yizhao Luan, Nan Tang, Jiaqi Yang, Congying Chen, Shuting Liu, Chichi Cheng, Yan Wang, Ya-nan Guo, Hongwei Wang, Mengqing Xiang, Rong Ju, Zhi Xie

**Author notes:** Equally contributed (Yizhao Luan, Nan Tang, Jiaqi Yang).

## Abstract

Human ribosomes, made of around 80 ribosomal proteins (RPs) and four ribosomal RNAs, have long been thought as uniform passive protein-making factories with little regulatory function. Recently, accumulating evidence showed heterogeneity of RP composition in ribosomes responsible for regulating gene expression in development and tumorigenesis. However, a comprehensive understanding of regulatory spectrum of RPs is still lacking. In this study, we conducted a systematic survey of regulatory specificity of human RPs on global gene expression. We quantified translational and transcriptional changes of gene expression upon deficiency of 75 RPs, including 44 from the large subunit (60S) and 31 from the small subunit (40S), by ribosomal profiling and RNA sequencing analysis. We showed that RP deficiency induced diverse expression changes, particularly at the translational level. RPs were subjected to co-translational regulation under ribosomal stress where deficiency of the 60S and the 40S RPs had opposite effects on the two subunits. RP deficiency perturbed expression of genes related to cell cycle, cellular metabolism, signal transduction and development. Deficiency of RPs from the 60S led to a greater repression effect on cell growth than that from the 40S by perturbing P53 signaling and cell cycle pathways. To demonstrate functional specificity of RPs, we showed that *RPS8* deficiency stimulated cellular apoptosis and *RPL13* or *RPL18* deficiency promoted cellular senescence. We also showed that *RPL11* and *RPL15* played important roles in retina development and angiogenesis, respectively. Overall, our study demonstrated a widespread regulatory role of RPs in controlling cellular activity, providing an important resource which can offer novel insights into ribosome regulation in human diseases and cancer.

## Introduction

Translation is a key step regulating gene expression. Depending on environmental, developmental and pathological conditions, translation is regulated by both *cis*-regulators, such as mRNA sequences and structures, and *trans*-regulators, such as eukaryotic initiation, elongation and termination factors [1]. Ribosomes are the main effector of the translational machinery to synthesize proteins. Although mammalian ribosomes are made of around 80 ribosomal proteins (RPs) and four ribosomal RNAs, ribosomes have long been thought as passive uniform molecular factories with little regulatory role [2].

Accumulating evidence in recent years suggested that developmental, pathological and stress conditions affected the composition of the ribosomes. Two paralogs of *RPL16* were mutual exclusively expressed in different organs of plants [3]. *RPL10A*, *RPL38*, *RPS7* and *RPS25* had varied levels in mouse embryonic stem cells [4]. In addition to normal tissues, RP composition heterogeneity was also found to be specifically associated with human cancers. For example, deletion of some RP genes was observed in about 43% TCGA cancer specimens [5]. Furthermore, many RPs exhibited strong dysregulation only in particular cancer types. For example, *RPL26L1* and *RPS27L* were exclusively up-regulated in breast and thyroid carcinomas, while expression of *RPL21* decreased in breast and uterine cancers [6].

Ribosomal heterogeneity can lead to functional specification of ribosomes, noted as “specialized ribosomes”, which preferentially translate a subset of transcripts [7]. The early demonstration of functional specification of RPs showed that ribosomes with *RPL38* deficiency impacted translation of a group of Hox mRNAs crucial for the formation of the mammalian body plan [8]. In yeast, *RACK1*-depletion repressed adaptive responses of yeast cells under amino acid deprivation condition [9]. *RPS26*-depletion ribosomes in plants preferentially translated mRNAs from selected stress response pathways during high-salt and high-pH stress [10].

Although these studies provided evidence for regulatory roles of several RPs, a systematic understanding of regulatory spectrum of RPs is unclear. In this study, we conducted a survey of regulatory specificity of human RPs on global gene expression. We characterized genome-wide gene expression changes at the translational and transcriptional levels after knocking-down 75 individual human RPs using ribosome profiling (Ribo-seq) and RNA sequencing (RNA-seq). Our results revealed that RP deficiency induced divergent gene expression changes, particularly at the translational level. RPs were subjected to co-translational regulation upon RP deficiency where deficiency of the large subunit (60S) and the small subunit (40S) had opposite effects on the two subunits. RP deficiency perturbed genes associated with a wide range of biological functions, including cell cycle, cellular metabolism, signal transduction and development. Deficiency of RPs from the 60S had more dramatic impacts on cell growth repression than that from the 40S by affecting P53 signaling and cell cycle pathways. To demonstrate functional specificity of RPs, we showed that *RPS8* deficiency stimulated cellular apoptosis while *RPL13* or *RPL18* deficiency promoted cellular senescence. We also showed specific regulatory roles of *RPL11* and *RPL15* in retina development and angiogenesis, respectively. Overall, our study demonstrated a widespread regulatory role of RPs in controlling specific cellular activity and provides an important resource and new biological insights into ribosome regulation.

## Results

### Overview of the transcriptome and translatome analysis

Aiming to gain a comprehensive understanding of RP regulation in human cells, we applied parallel RNA-seq and Ribo-seq to human A549 cells with each individual depletion of 78 RPs via specific siRNAs. At least two siRNAs for each RP were designed to avoid off-targeting (Table S1). The knocking-down efficiencies were tested and confirmed via RT-PCR (Table S2). In total, we prepared 105 Ribo-seq libraries, with 105 paired RNA-seq libraries, for 78 human RPs as well as 8 control (untreated) samples, where biological replicates were performed for 16 RPs and for 3 control samples. Ribo-seq and RNA-seq datasets of 75 RPs passed stringent quality controls and were kept for further analyses (Materials and Methods) [11]. In total, Ribo-seq and RNA-seq experiments generated around 14 and 3.6 billion clean reads aligned to the human reference genome, with an average of around 33 and 32 million uniquely mapped reads per library (Table S3).

For each library, ribosome-protected fragments (RPFs) showed expected size distributions mainly ranging from 28 to 32 nucleotides (nt) (Figure 1A). While RNA-seq libraries were evenly mapped to transcripts along 5’ to 3’ orientation (Figure S1A), RPFs presented characteristic 5’ ramps where ribosome footprints declined along gene-body (Figure S1B) [12]. Both Ribo-seq and RNA-seq reads showed significantly higher density on the coding sequence (CDS) (t-test, all *P* < 2.2e-16) (Figure S1C, D). The relative percentages of RPF distributed on CDS, 5’ untranslated region (UTR) and 3’ UTR were 97.7%, 2.1% and 0.2%, respectively, which showed strong preference to the CDS and depletion in 3’ UTR compared to RNA-seq reads (87.6% on CDS, 1.0% on 5’ UTR and 11.4% on 3’ UTR) (Figure 1B, C), displaying a typical read distribution on genomic regions for RPF and RNA-seq [13]. As expected, RPF mapped to the CDS showed a clear 3-nucleictide periodicity, with ∼58% of reads on the first reading frame in average (Figure 1D-F). In contrast, RNA-seq reads showed no frame preference (Figure S1E). The Pearson correlation coefficients (PCC) of biological replicates ranged from 0.9760 to 0.9962 for the Ribo-seq (a median of 0.9936) and from 0.9576 to 0.9929 for the RNA-seq (a median of 0.9884) experiments, illustrating high reproducibility of our experiments (Figure 1G; S1F, G). The control experiments also showed high correlation with the public A549 Ribo-seq data (PCC R=0.81, *P* < 2.2e-16) and RNA-seq data (PCC R=0.79, *P* < 2.2e-16) (Figure S1H, I; Materials and Methods). Taken together, these results suggested that our data were of high-quality and highly reproduceable.

**Figure 1.**
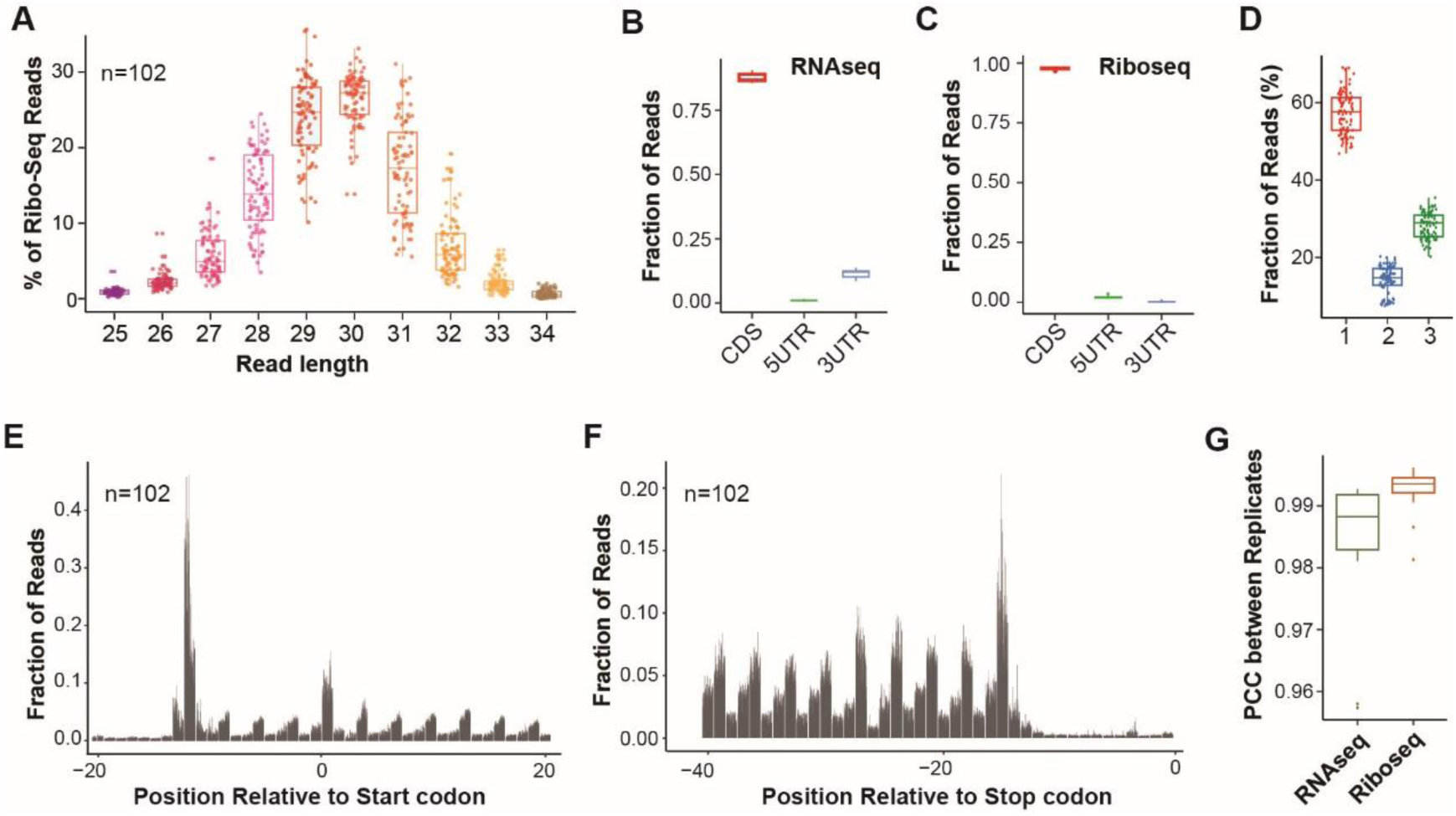
Overview of the transcriptome and translatome datasets. (A) Length of RPFs in Ribo-seq libraries for all samples. Each point indicates one library. (B, C) The relative fraction of reads mapped to the CDS, 5’ UTR and 3’ UTR of annotated transcripts in RNA-seq (B) and Ribo-seq (C) libraries. When genome features were overlapped, CDS exons were prioritized over UTR exons. (D) Percentages of RPFs mapped to reading frames by combining all Ribo-seq libraries. Each point indicates one library. (E, F) Fractions of reads assigned to each nucleotide around the start codons (E) or stop codons (F) for all libraries in Ribo-seq data. Each bar indicates one sample. P-site was used to assign the short read to transcript location. (G) Pearson correlation coefficients between replicates in RNA-seq and Ribo-seq datasets. Log2 RPKM values were used in the correlation analysis.

### Landscape of gene expression changes

Comparing RP-deficient cells with the control cells, we identified 9,264 differentially translated genes (DTGs) by Ribo-seq and 6,743 differentially expressed genes (DEGs) by RNA-seq for 75 RPs (Materials and Methods). Each RP deficiency impacted distinct genes at both the transcriptional and translational levels (Figure S2A, B), where DTG number ranged from 317 to 2,631, with a median of 1,299, and DEG number ranged from 97 to 1,676, with a median of 476 (Figure 2A). 73% of DEGs and 51% of DTGs were regulated by no more than five RPs while only 7.6% DEGs and 18.36% DTGs were regulated by more than 20 RPs, a behavior following a power-law distribution (Figure 2B). We observed significantly more DTGs than DEGs for 72 RPs (Wilcoxon test, *P* < 2.2e-16; Figure 2C, S2C) and larger changes in magnitude for the DTGs than the DEGs, measured by the difference of 97.5% and 2.5% quantiles of the fold changes (Wilcoxon test, *P* < 2.2e-16; Figure 2D). These together indicated that RP deficiency caused more drastic changes at the translational level than the transcriptional level.

**Figure 2.**
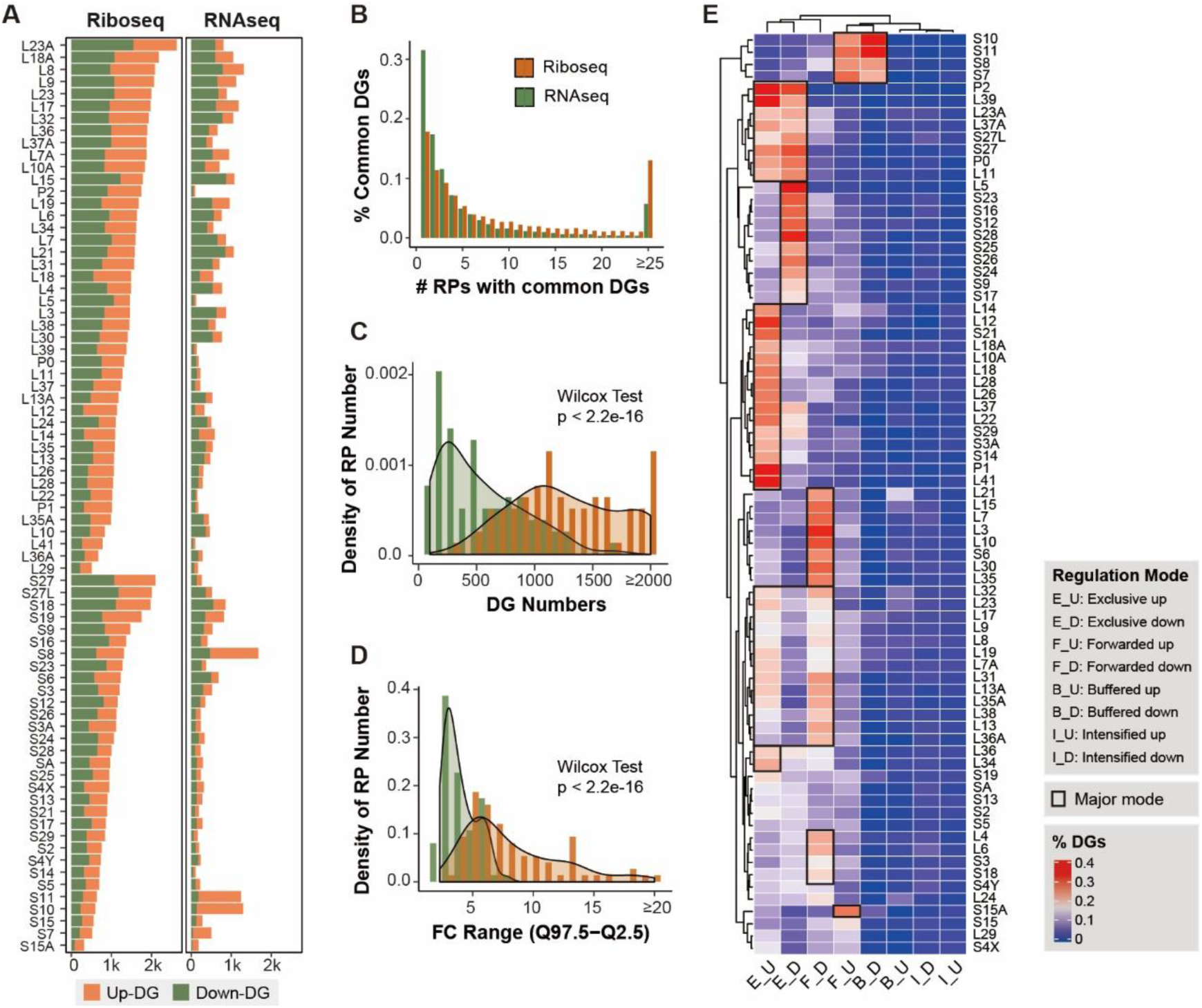
Diversity of gene expression changes upon RP knockdown. (A) Numbers of up- and down-regulated differential genes in RNA-seq and Ribo-seq for all RPs. RPs are ranked by DTG numbers in Ribo-seq. (B) Frequency of common DEGs or DTGs between RPs in RNA-seq and Ribo-seq. (C) Distribution of DEG or DTG numbers for all the RPs. The distributions in RNA-seq and Ribo-seq were compared by Wilcoxon sign-rank tests. (D) Ranges of global gene fold changes for all RPs. The ranges in RNA-seq and Ribo-seq were compared by Wilcoxon sign-rank tests. (E) Percentages of genes under different gene expression regulation modes after knockdown of each RP. RPs were grouped by hierarchical clustering analysis of the regulation profiles. Rectangles indicates the primary regulation mode for RP groups.

Comparing DEGs and DTGs for each RP showed that the percentage of overlapped genes, measured by Jaccard index, ranged from 2.72% to 46.50%, with a median of 20.90%, indicating that RP deficiency induced a large degree of uncoupled gene expression changes transcriptionally and translationally. We next accounted for the transcriptional and translational contributions to gene expression changes upon RP deficiency. The differential genes identified in RNA-seq and Ribo-seq for each RP were categorized into four regulatory modes, including intensified, buffered, exclusive, or forwarded regulation, based on their changes of RPF reads, mRNA reads and translational efficiency (TE) (Figure S2D). Specifically, the forwarded genes have RPF changes that are explained by the mRNA changes. The exclusive genes have changes in TE without mRNA changes. The buffered and intensified genes have changes in TE that offsets or amplifies the mRNA changes, respectively. Out of 75 RPs, 66 can be categorized into one or two regulation modes (Figure 2E). Most of deficiency of RPs (35) induced exclusive regulation while 13 showed forwarded regulation. In addition, 13 showed both exclusive and forwarded regulation. These results indicated that deficiency of individual RPs mainly regulated expression of specific subsets of genes at the translational level. Further functional analysis showed that the gene expression changes under the same regulation mode by different RPs enriched in specific functions (Figure S2E-K). For example, down-regulated genes by translational exclusive regulation of *RPS12* were related to transferase activity while that of *RPS26* were related to aminoglycan metabolic process (Figure S2E).

### Co-regulation of 60S and 40S RPs after RP deficiency

Previous studies showed that individual RP deficiency could result in dysregulation of other RPs, further leading to imbalance in stability and accumulation of ribosomal subunits [14, 15]. Our dataset offers an opportunity for a systematic investigation of co-regulation of RPs upon RP disruption in human cells. Substantial downregulations of each RP targeted by specific siRNAs were observed at both the transcriptional and translational levels (the diagonal line in Figures 3A and S3A), confirming high knockdown efficiencies of siRNA transfection in our dataset. A distinct pattern was observed at the translational level where knocking-down 60S RPs repressed expression of RPs of both subunits while knocking-down 40S RPs resulted in elevation of RPs of both subunits. At the transcriptional level, knocking-down RPs had smaller effects on the expression changes of the other RPs (t-test, *P* = 4.41e-15; Figure S3B). Interestingly, we also observed opposite effects of perturbation of RPs from the 60S and the 40S, where knocking-down 60S RPs slightly activated expression of both subunits while knocking-down 40S RPs slightly repressed expression of both subunits (Figure S3A). Further TE analyses demonstrated that knocking-down 60S RP led to global TE downregulation whereas knocking-down 40S RP resulted in global TE upregulation of RPs (Figure 3B).

**Figure 3.**
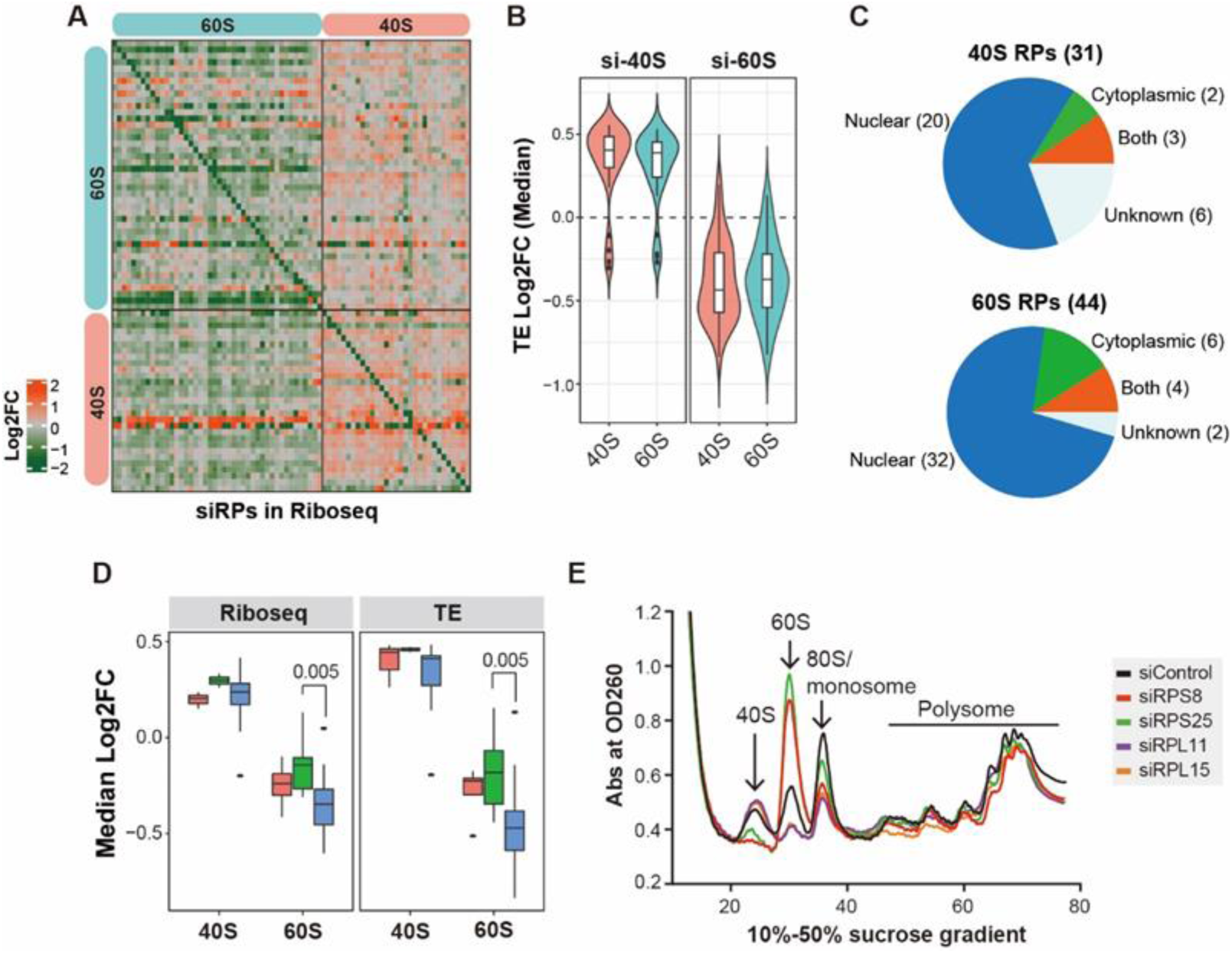
Co-regulation of 60S and 40S RPs. (A) The expression changes of RPs (rows) after knockdown of individual RPs (columns) in Ribo-seq dataset. Log2 fold change values are shown. RPs for rows and columns according to ribosomal subunits as indicated by colored sidebars: cyan color indicates 60S RPs, orange color indicates 40S RPs. (B) Comparison of global changes in TE of 40S and 60S RPs after knockdown of RPs from 40S (left) or 60S (right). Median TE values of the remaining RPs were used for each RP targeted by siRNAs. (C) Numbers of RPs annotated to cellular locations where ribosomal assembly occurs. (D) Comparison of expression changes at the translatome level (left) and TE changes (right) of the remaining RPs after knockdown of individual RPs within different stages of ribosomal subunit assembly. Wilcoxon tests were performed to compare the difference between groups for each subunit. (E) Polysome profiling by sucrose-gradient-based centrifugation showing the changes in abundance of ribosomal components (40S, 60S, 80S monosome and polysomes) in A549 cells treated by specific siRNAs targeting *RPS8*, *RPS25*, *RPL11* or *RPL15*.

Because RPs enter the assembly pathway at different time points where ribosome assembly first occurs in the nucleus and continues in cytoplasm [16], we suspected that knockdown of the RPs that enter the assembly pathway earlier might have larger impact on RP translation than the other RPs. To test the hypothesis, we annotated the RPs based on the locations they assemble, that is nucleus, cytoplasm, or both (Figure 3C). We found that knocking-down 60S RPs assembled in the nucleus reduced higher degree of the expression of RPs at the translational level and TE level in comparison with those assembled in the cytoplasm (Wilcoxon test, both *P* = 0.005; Figure 3D), suggesting that deficiency of 60S RPs entering the assembly pathway earlier had greater effect on the translational repression of RPs than those entering later. In contrast, we found no such assembly stage dependent regulation upon knockdown of 40S RPs (Wilcoxon test for RPF: nuclear vs cytoplasmic, *P* = 0.3117; nuclear vs both, *P* = 0.4574; Wilcoxon test for TE: nuclear vs cytoplasmic, *P* = 0.1039; nuclear vs both, *P* = 0.4043).

We also investigated the impact of deficiency of 60S or 40S RPs to ribosome assembly. Polysome profiling by sucrose-gradient-based centrifugation can assess relative abundance of 40S and 60S subunits, where the peak at the point in the gradient of corresponding subunit reflects the abundance of that subunit. Polysome profiles of four RPs were analyzed, two RPs from 60S (*RPL11* and *RPL15*) and two from 40S (*RPS8* and *RPS25*). On one hand, knockdown of *RPL11* or *RPL15* decreased the abundance of 60S but increased that of 40S (Figure 3E). On the other hand, knockdown of *RPS8* or *RPS25* increased 60S and decreased 40S (Figure 3E). Taken together, our analysis demonstrated that deficiency of 40S or 60S RPs had different effects on 60S and 40S subunits, at the transcriptional and translational levels, as well as at the ribosome assembly stage.

### Global functional characterization of RPs

We next functionally annotated RPs based on the gene ontology (GO) analysis. In total, we obtained enriched GO terms for 64 RPs (Table S4). The enriched GO terms can be grouped into eight major classes, mainly including cell cycle, organelle organization, nucleotide and other macro-molecular metabolism, signal transduction, cell response, cell and tissue development, and transport (Figure 4A). We knock-downed some RPs using another cell line, HCT116, and the enriched terms, such as neurogenesis in *RPL11*, cell cycles in *RPS6* and *PRS8*, could also been confirmed (Figure S4A-C). Based on extensive literature search, we found that many previously known functional terms of RPs could be recovered in our study, such as *RPS6* and *RPL15* in cell cycle, *RPL17* in sex differentiation, *RPS6* in immune system development and *RPL15* in Diamond-blackfan Anemia (DBA) disease (Figure 4B; Table S5). These together indicated that our datasets could be used to infer functions of RPs.

**Figure 4.**
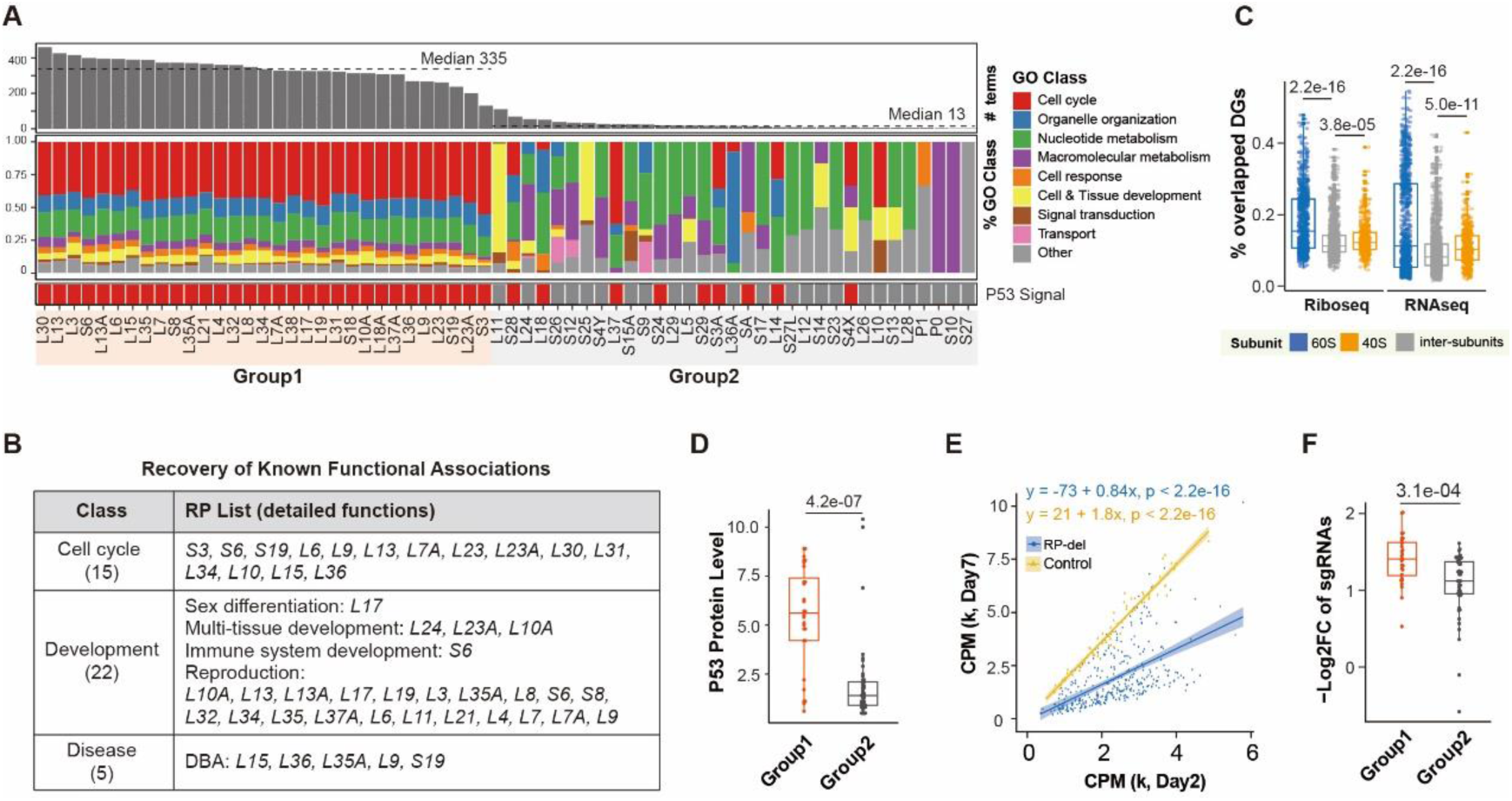
Functional characterization of RPs. (A) 64 RPs are ranked according to the number of significantly enriched GO BP terms. Annotation bar at the bottom indicates the enrichment of P53 signaling pathway after RP knockdown; Middle heatmap shows the fraction of GO classes; Bar plot at the top shows total number of enriched GO BP terms for each RP. (B) Summary of the recovered known functional associations of RPs. Details can be found in Table S5. (C) Percentages of overlapped DEGs or DTGs after knockdown of 40S or 60S RPs. T-tests were performed. (D) Comparison of P53 protein levels after knockdown of Group1 and Group2 RPs. P53 protein levels were extracted from [11]. (E) Comparison of sgRNA expression levels at Day 2 and Day 7 after transfection. Linear regression models were estimated and tested for sgRNAs with RP targets and without targets. The estimated slope values indicate cell growth rate over time: slope>1 indicates positive cell growth, slope < 1 indicates repressed cell growth. (F) Comparison of expression changes over time between sgRNAs targeting RPs from Group1 and that from Group2. T-test was performed.

Based on the number of enriched GO terms, RPs can be categorized into three functional groups: 31 RPs (Group1) had a large number of enriched GO terms (a median of 335), indicating their extensive functional associations, while 33 RPs (Group2) enriched a small number of GO terms (a median of 13), indicating their selective functions (Figure 4A). In addition, 11 RPs did not have enriched GO terms. Notably, Group1 was significantly enriched by 60S RPs (26 out of 31 RPs, Fisher’s exact test, *P* = 0.0234), suggesting that deficiency of 60S RPs had generally a larger impact on cellular functions than 40S RPs. This was consistent with a previous study showing that knockdown of 60S RPs induced larger nuclear disruption than 40S RPs [11]. Although deficiency of 60S and 40S induced similar numbers of DEGs (t-test, *P* = 0.10), 60S RP deficiency led to significantly more DTGs than that by 40S RPs (t-test, *P* = 0.018; Figure S4D). Moreover, 60S RP deficiency usually caused greater magnitude of changes for DEGs and DTGs (t-test, *P* < 0.05; Figure S4D). As expected, RPs from the same subunit shared significantly more DEGs or DTGs than the RPs from different subunits (Wilcoxon test, *P* < 0.05; Figure 4C), indicating that RPs from the same subunit tended to perturb expression of similar genes.

It is notable that all the RPs in Group1 induced changes in P53 signaling and cell cycle pathways upon knockdown. Moreover, deficiency of these RPs increased P53 protein levels more significantly than that in Group2 (t-test, *P* = 4.2e-07; Figure 4D). We further experimentally test whether the disruption of the Group1 RPs had a larger impact on the cell growth comparing with that of Group2 by CRISPR-Cas9 experiments. We knocked-out individual RPs with single guide RNAs (sgRNAs) (Materials and Methods) [17] and found that cells treated by RP-targeting sgRNAs were repressed (F-test, slope=0.84, *P* < 2.2e-16), while cells treated by non-targeting sgRNAs grew over time (F-test, slope=1.8, *P* < 2.2e-16; Figure 4E). Further comparing the foldchanges of RP-targeting sgRNAs revealed that disruption of the Group1 RPs significantly repressed cell growth more than that by the Group2 RPs (t-test, *P* = 3.1e-04; Figure 4F), illustrating the larger repression on cell growth by the deficiency of the Group1 RPs than the Group2.

### RPs regulate cell cycle and promote divergent cell fates

Our data showed that deficiency of many RPs perturbed cell cycle related pathways. We next selected five RPs (*RPS8*, *RPL13*, *RPL18*, *RPL22*, and *RPL29*) to validate their effects on cell cycle progression and cell proliferation upon knockdown. Among them, *RPS8*, *RPL13* and *RPL18* were shown to be associated with cell cycle in our datasets while *RPL22* and *RPL29* were not. The functional association of *RPS8* and *RPL18* with cell cycle had not been previously reported.

After these RPs were efficiently knocked-down (Figure S5A), flow cytometry analysis revealed that knockdown of *RPS8, RPL13* or *RPL18* significantly increased the G1 phase cell populations and decreased the S phase cells at 24h post treatment (t-test, *P* < 0.001 for all the three RPs), suggesting a possible G1 arrest. On the other hand, knockdown of *RPL22* or *RPL29* did not change the phase distributions in comparison to the control cells (t-test, *P* > 0.05; Figure 5A, S5B). Consistently, knockdown of *RPS8, RPL13* or *RPL18* significantly inhibited cell proliferation in MTT assays at 72h after siRNA transfection with various degrees (t-test, *P* < 0.01 for all the three RPs, Figure 5B). In contrast, *RPL22* or *RPL29* had little impact on cell proliferation and no difference was observed in the cells lacking *RPL22* or *RPL29* at 72h post-transfection in comparison with the control cells (t-test, *P* > 0.05).

**Figure 5.**
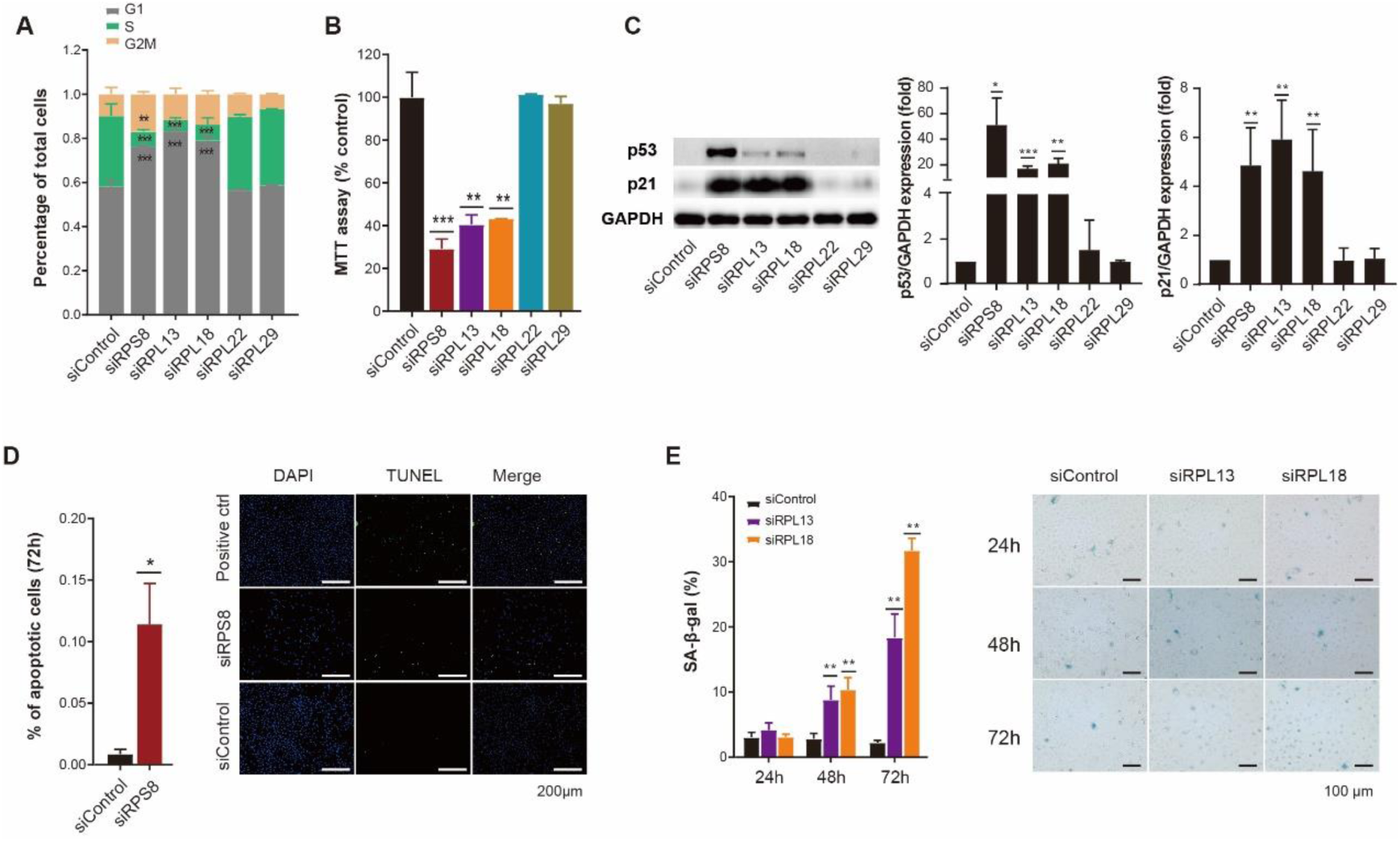
RP deficiency promoted divergent cell fates after cell cycle arrest. (A) Percentages of cells within different cell cycle stages (G1, S, and G2/M) by flow cytometry experiments on A549 cells at 24hr after knockdown of indicated RPs. Three replicates were used in t-tests. (**), p<0.01; (***), p<0.001. (B) Changes of cell viability by MTT assays on A549 cells at 24hr after knockdown of indicated RPs. Three replicates were used in t-tests. (**), p<0.01; (***), p<0.001. (C) Representative of western blotting assays (left panel) and quantification of p53 (middle panel) or p21 (right panel) protein levels at 24hr after knockdown of indicated RPs (n=2 for p53 or 3 for p21 tests). T-tests were used. (*), p<0.05; (**), p<0.01; (***), p<0.001. (D) Bar plots (left panel) showing the percentage of TUNEL+ cells at 72hr after knockdown of *RPS8*. Representative of TUNEL staining assays (right panel) for testing apoptosis *in situ* in A549 cells at 72hr after knockdown of *RPS8*. Three replicates were used in t-test. (*), p<0.05. (E) Bar plots (left panel) showing the percentages of *β*-gal-positive cells at 24hr, 48hr and 72hr after knockdown of *RPL13* or *RPL18*. Representative of *β*-gal staining assays (right panel) for testing senescence in A549 cells at 24hr, 48hr and 72hr after knockdown of *RPL13* or *RPL18*. Three replicates were used in t-tests. (**), p<0.01.

To explore the underlying mechanisms, the protein levels of p53 and p21, two well-known cell cycle regulators [18], were examined by western blotting assays. As expected, the protein levels of p53 and p21 in the cells lacking *RPL22* or *RPL29* showed no change compared to the control cells. On the contrary, the levels of p53 and p21 elevated in the cells lacking *RPS8, RPL13* or *RPL18* (t-test, *P* < 0.05 for all three RPs (p53), *P* < 0.01 for all three RPs (p21); Figure 5C). Moreover, although knockdown of three RPs all led to increased p21 to the similar degrees, the protein levels of p53 appeared to be different, suggesting that the cell cycle arrest mediated by RP knockdown could go through different mechanisms: *RPS8* deletion arrested cell cycle via activation of the p53/p21 signaling while *RPL13* or *RPL18* deficiency triggered p21 expression likely due to accumulation of p53 partially.

It is well known that increased p21 protein levels would lead to different cellular consequences, senescence or apoptosis [19]. To further explore possible cell fates by knocking-down *RPS8*, *RPL13* or *RPL18*, we performed TUNEL and SA-*β*-gal staining assays. Interestingly, in comparison with the control cells, we observed significantly more TUNEL-positive nuclei lacking *RPS8* at 72h after siRNA transfection (t-test, *P* < 0.05), suggesting stimulated cellular apoptosis of these cells (Figure 5D, S5C). In contrast, we observed gradually increased *β*-gal-positive cells over time from 48h to 72h after knockdown of *RPL13* (t-test, *P* < 0.01) or *RPL18* (t-test, *P* < 0.01), suggesting increased senescent events induced by deficiency of *RPL13* or *RPL18* (Figure 5E, S5D). Collectively, our results showed that different RPs would influence cell cycle and cell fate differently through different mechanisms.

### RPs regulate development

Previous studies showed ribosomal heterogeneity might impact development by regulating specific mRNAs [4, 8, 20]. The GO analysis based on our current datasets indicated that RP deficiency changed expression of genes associated with a wide range of tissue or organ development, including cardiovascular, respiratory, digestive, urogenital, reproductive and nervous systems (Figure S6A). Of note, some development relevant functions of RPs had been previously reported. For example, our data showed that *RPL17* deficiency impacted genes related to sex differentiation, consistent with the observation that *RPL17* expressed differently in male and female brains of developing zebra finches [21]. Deficiency of *RPL24*, *RPL23A* or *RPL10A* affected genes involved in embryonic development, consistent with the phenotypes of their mutant in developing embryos of zebrafish [22, 23] or mouse[24]. On the other hand, we also identified many RPs showing specifically regulatory potential in the development of tissues that had not been reported, among which were *RPL11*, its deficiency associated with neurogenesis, and *RPL15,* its deficiency associated with angiogenesis (Figure 6A).

**Figure 6.**
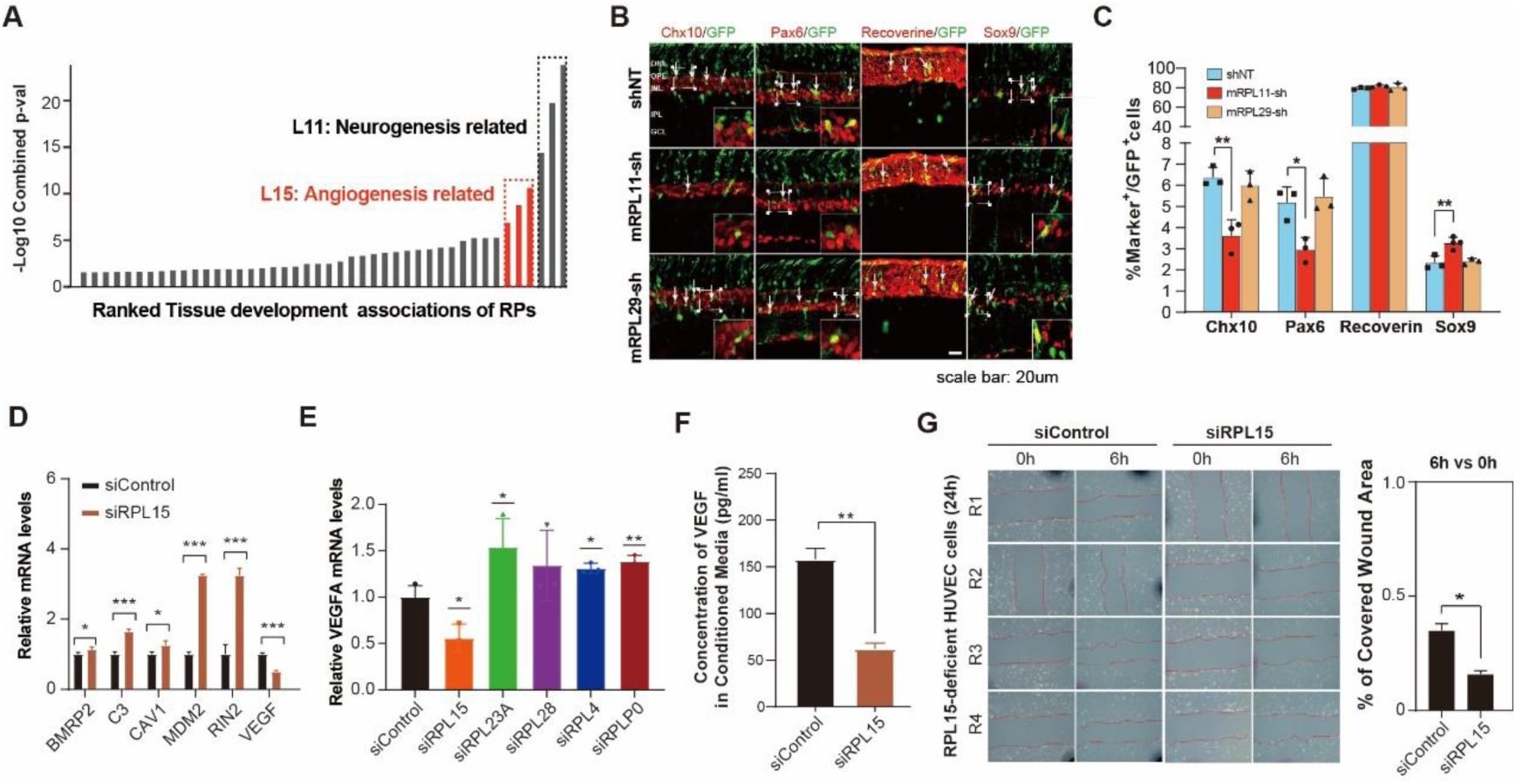
RP deficiency regulates tissue development. (A) The ranked tissue development terms affected by RP knockdown according to the significance. Tissue development terms were defined according to GO structure. The p-values for all subentries belonging to the term of interest were combined with Fisher’s method. Grey bars indicate GO terms enriched by down-DTGs; Red bars indicate GO terms enriched by up-DTGs. (B, C) Analysis of developing retinal cells after *in vivo* conditional knockdown of *RPL11*. (B) P0 retinas co-electroporated RP-targeting shRNAs or pU6 plasmid with the pCIG vector were collected at P12, and their sections were double-immunostained with an anti-GFP antibody and antibodies against Chx10, Pax6, Sox9 or Recoverine. Arrows point to representative colocalized cells. (C) The numbers of specific cell types and statistical testing results between groups. Two or three replicates were used in t-tests. (*), p<0.05; (**), p<0.01. (D) The relative mRNA levels by qPCR of representative genes in *RPL15*-deficient A549 cells. Three replicates were used in t-tests. (*), p<0.05; (**), p<0.01; (***), p<0.001. (E) The relative mRNA levels of VEGFA in the control HUVEC and the HUVEC upon knockdown of indicated RPs. Three replicates were used in t-tests. (*), p<0.05. (F) ELISA analysis showing the concentration of VEGF proteins in conditioned media from the control A549 cells and *RPL15*-deficient A549 cells. T-test was used. (**), p<0.01. (G) Images of the control HUVEC and *RPL15*-deficient HUVEC 0hr and 6hr after a scratch was introduced in the monolayer with a pipette tip (left panel). The percentage of open wound area for each replicate at 6hr over that at 0hr was estimated and compared between *RPL15*-deficient cells and the control cells (right panel). Four replicates were used in t-test. (*), p<0.05.

To validate regulation of neurogenesis by *RPL11*, we co-electroporated RP-targeting small hairpin RNAs (shRNAs) into mouse retinas at P0 (postnatal day 0) and collected them at P12. The proportions of GFP-positive progeny distributed in different retinal cell layers at P12 were quantified. We observed significantly less Pax6-positive amacrine cells (t-test, *P* < 0.05) and Chx10-positive bipolar cells (t-test, *P* < 0.01) in retina of *RPL11*-knockdown mice than that in the control mice. By contrast, expression of *RPL11* shRNA increased the percentage of Sox9-positive Müller cells (t-test, *P* < 0.01) and had no effect on the differentiation of Recoverin-positive photoreceptors (Figure 6B, C). We also tested *RPL29* which had no effect on neurogenesis from our analysis as a negative control. As expected, we did not observe any changes in all the cell subtypes from retinas of *RPL29*-knockdown mice (Figures 6B, C). These results indicated that *RPL11* deficiency would inhibit retinal cell differentiation into amacrine and bipolar cells but promoted cell differentiation to Muller cells.

To investigate the potential involvement of *RPL15* in angiogenesis, we firstly validated the mRNA expression changes of several angiogenesis-associated genes, including *BMPR2*, *C3*, *CAV1*, *MDM2*, *RIN2* and *VEGFA*, by real-time PCR. As expected, the mRNA level of *BMPR2*, *C3*, *CAV1*, *MDM2* and *RIN2* increased whereas mRNA of *VEGFA* decreased upon *RPL15* knockdown (t-test, all *P* < 0.05; Figure 6D), consistent with our sequencing data. *VEGFA* is known to be the most important proangiogenic factor. It is secreted by many types of cells including diverse cancer cells and endothelial cells to promote blood vessel growth [25]. We also confirmed that the mRNA level of *VEGFA* was downregulated in the human umbilical vein endothelial cells (HUVECs) when *RPL15* was knocked down (t-test, *P* < 0.05; Figure 6E). In addition, the VEGFA proteins secreted by *RPL15*-deficient cells detected by ELISA analysis was significantly decreased as well (t-test, *P* < 0.01; Figure 6F). To further determine the biological impact of *RPL15* deletion, we knocked down this gene in HUVECs and performed the migration assay. The result showed that *RPL15*-deficiency inhibited cell migration in comparison with the control siRNA treatment (t-test, *P* < 0.05; Figure 6G). Since VEGFA produced by tumor cells can promote angiogenesis in order to support their growth [25], we also carried out experiments to imitate such a situation. We treated HUVECs with the conditioned medium (CM) from the *RPL15*-deficient A549 cells and examined how the CM impacted cell migration and tube formation. We found no differences in cell migration and tube formation of the HUVECs treated by the CMs from the *RPL15*-deficient A549 cells and the control cells (Figure S6B-D), suggesting that the effects of decreased *VEGFA* might be offset by other proangiogenic factors increased by *RPL15* deficiency. These results together demonstrated that *RPL15* deficiency impacted *VEGFA* expression and its potential role in angiogenesis via affecting migration of endothelial cells.

## Discussion

Our study represents the first systematic effort to study the regulatory roles of RPs in mammalian cells. Although ribosome heterogeneity is evident and functional specification of RPs has been suggested, we still have limited understanding of specific functions by which RPs might act, and even more rare is the understanding in physiological consequence of the functional specialization. In this study, we integrated Ribo-seq and RNA-seq to systematically investigate the genome-wide impacts of RP deficiency on gene translation and transcription. Our study expands the knowledge of regulatory roles of ribosomes and provides novel insights into the complex functions of ribosomes and translational regulation.

While it has been known that knockdown of RPs that have little effects on overall translation, has minor effects on transcriptome [13], our data demonstrated that RP deficiency generally has larger effects on translatome than transcriptome, independent of whether the depleted RP can affect global translation. And the diversity in molecular changes and functional associations after RP deficiency can be enlarged mainly by translational regulation.

Of note, our data demonstrated that the RPs from the 60S subunit were generally different from that from the 40S subunit in many aspects. (1) 60S RP deficiency usually resulted in more gene expression changes and more frequent elevation of genes downstream of p53 signaling. Moreover, genomic depletion of RPs from 60S repressed cell growth more. One possible mechanism for this difference involved the perturbation of highly conservative feedback loop between MDM2 and P53 [26]. Several RPs, such as *RPL5* [27], *RPL11* [28] and *RPL23* [29], can directly bind to MDM2, thus promote the stability of p53 protein and activate p53 signaling pathway. It has been known that depletion of 60S RPs could increase p53 protein level more significantly [11]. (2) Deficiency of 60S and 40S RPs induced distinct regulations on ribosomal subunits at the transcriptional, translational and assembly stages. While the regulation of the remaining RPs after individual RP deficiency are important for maintaining cell homeostasis, the regulatory effects have not been determined. Previous studies showed opposite effects. While genomic depletion of individual RP from yeast genome mainly upregulated genes involved in ribosome biogenesis at the translation level [30], deficiency of selected RPs in hematopoietic stem and progenitor cells mainly translationally repressed RP genes [31]. Our data demonstrated that 60S and 40S RPs were subjected to different translational effects after RP deficiency: 60S RP knockdown translationally repressed RPs of two subunits and 40S RP knockdown translationally upregulated all the other Rps (Figure 3A, B). Our further polysome profiling analysis, on one hand, demonstrated expected changes of abundance of the ribosomal subunit to which the targeted RP belong. On the other hand, the complementary subunit showed increased abundance after the targeted RP was knockdown. Increased 40S after 60S RP knockdown was also observed in other studies, probably due to the accumulation of 40S that failed to combine with 60S to initiate translation because of blocked 60S assembly; Increased 60S after 40S RP knockdown was consistent with the notion that 60S assembly would not be affected by blocked 40S assembly [14]. These results also agreed with the mechanism in mammalian cells that ribosomal proteins levels are never rate limiting for the efficient assembly of ribosome subunits if needed [32]. 3) Another possible mechanism for this difference involved the *cis*-regulatory elements in mRNA targets, particularly in the 5’ UTR regions. It is known that 40S subunits are firstly recruited to the mRNA-cap and 60S are then combined once the start codons occur [33]. Based on this model, researchers have hypothesized that translation of mRNAs containing elements blocking 40S scanning would be sensitive to 40S subunit concentration while translation of mRNAs with a poor Kozak context would be sensitive to 60S subunit concentration [7]. We therefore explored the mRNA elements possibly involved in translational regulation after RP deficiency and found that knockdown of RPs from the 40S subunit usually downregulated TE of genes with higher GC content in 5’ UTR regions while knockdown of RPs from the 60S subunit usually upregulated TE of genes with strong Kozak sequence with cytosines (C) at −1, −2 positions and with guanine (G) at −3 position and downregulated TE of genes without these signals (Figure S7). Further studies will be required to explore the complex interaction between multiple *cis*-regulatory elements.

Our data suggests many important functional preferences of RPs. We showed that our findings were unlikely to be affected by cell-type-specific gene expression by recovering many known functional associations of RPs and by computationally confirming a low overlap between A549 cell-specific genes and our defined differential genes (Figure S8). To validate the functional relevance, we explored and observed similar gene expression changes in other cells such as HCT116 and HUVEC. We further presented several examples to demonstrate the value of our datasets in exploring novel regulatory roles and functions of RPs. We for the first time showed the repression effects of *RPS8, RPL13* or *RPL18* deficiency on cell cycle progress in human cancer cells, suggesting their anti-cancer potential because targeting cell cycle is considered to be an effective way for tumor suppression [34]. We showed novel physiological function of *RPL11* where *in vivo* knockdown of *RPL11* induced disordered neuronal differentiation in retinas. This extends our knowledge of regulatory functions of *RPL11,* which was thought to be involved in DBA, where previous studies showed that partial loss of *RPL11* in adult mice can result in DBA-mimic phenotypes and cancer predisposition to lymphomagenesis [35]. While there is opinion that neurodevelopment appears to be relatively resistant to partial impairments of ribosomal biogenesis that are sufficient to produce tissue/cell linage-specific phenotypes[36], our data directly showed that it’s not true for all RPs. Finally, we provided evidence of novel role of *RPL15* in angiogenesis. We showed that *RPL15* deficiency preferred to affect angiogenesis factors including *VEGFA* and found significant repression effect in cell migration of *RPL15*-deficient endothelial cells. While *RPL15* over-expression has been revealed to be associated with breast cancer metastasis [20], our data suggested that *RPL15* can also selectively regulate *VEGFA* to mediate cancer progression related cellular functions. Further studies will be required to test the functional significance of associations between RPs and other angiogenesis factors in diverse human cancers.

Overall, our study demonstrated a widespread regulatory role of RPs in controlling cellular activity. Our datasets provided an important resource which offered novel insights into ribosome regulation in human diseases and cancer.

## Materials and Methods

### Cell lines

A549 cells and Neuro-2a cells were obtained from American Type Culture Collection (ATCC). A549 cells were cultured in Dulbecco’s modified Eagle’s medium (DMEM) with 10% fetal bovine serum (FBS) (Gibco, 10270-106) and antibiotics (100μg/ml penicillin and 50μg/ml streptomycin sulfate) (Gibco, 15140122). Neuro-2a cells were maintained in modified Eagle’s medium (MEM) with 10% FBS, antibiotics (100μg/ml penicillin and 50μg/ml streptomycin sulfate) and 1% NEAA (Gibco, 11140050). HUVECs were bought from ScienCell Research Laboratories (ScienCell, 8000) and cultured in EGM-2 medium (Lonza, CC-3162). All cell lines were genotyped to confirm their identity at Genewiz. Cells were incubated at 37℃ with 5% CO_2,_ and tested routinely for Mycoplasma contamination.

### siRNA transfection

A549 cells were transfected for 24h with RP-targeting or non-targeting (control) siRNAs at a final concentration of 40nM using Lipofectamine RNAiMAX reagent (Invitrogen, catalog no.13778150) following the manufacturer’s protocol.

### Animals

All experiments on animals were performed according to the IACUC (Institutional Animal Care and Use Committee) standards, and approved by Zhongshan Ophthalmic Center, Sun Yat-sen University. The CD1 mice were purchased from the Vital River Laboratories (Beijing, China).

### Quantitative real-time PCR (qRT–PCR)

Total RNA was isolated with TRIzol (Invitrogen,15596018) following the manufactory protocol. cDNA was synthesized from 1µg total RNA using HiScript II Q RT SuperMix for qPCR (Vazyme, R223-01). Real-time PCR reactions were performed using iTaq™ Universal SYBR Green Supermix (Bio-rad,1725121) and CFX96 system (Bio-rad). Each reaction was performed in triplicate. The primers are shown in Table S6. The mRNA fold changes were calculated by the ΔΔ*Ct* method using the housekeeping gene GAPDH or *β*-actin as internal control and normalized to the experimental control.

### Library preparation for RNA-seq and Ribo-seq

A549 cells were harvested 5 minutes (min) after treatment with 100 μg/ml cycloheximide (CHX) (Sigma, C4859) in DMEM, washed twice in PBS with 100 μg/ml CHX. Samples were lysed with 1 ml mammalian lysis buffer containing 200 µl 5x Mammalian Polysome Buffer (Epicentre, RPHMR12126), 100 µl 10% Triton X-100, 10 µl 100 mM DTT, 12.5 µl Turbo DNase (Invitrogen, AM2239), 1 µl 100 mg/ml CHX, and 675.5 µl nuclease-free water. After incubation for 20min on ice, lysates were centrifuged at 12,000rpm, 4°C for 10min. For each sample, lysate was divided into two aliquots (about 600μL for Ribo-seq,150μL for RNA-seq). For the 600-μl aliquots lysates, 6 units of ARTseq Nuclease was added to each A260 lysate, and the mixtures were incubated for 60min at room temperature with rotation. Nuclease digestion was stopped by 10μL SUPERase·In RNase Inhibitor (Ambion, AM2696). Lysates were applied to MicroSpin S-400 HR spin columns (GE Healthcare, 27-5140-01). Total RNA was purified with Zymo RNA Clean & Concentrator-25 kit (Zymo Research). The rRNA was depleted with Ribo-Zero Gold rRNA Removal Kit (Illumina, MRZG126). The rRNA-depleted samples were run on a 10% TBE-Urea polyacrylamide gel. Ribosome-protected fragments between 28-nt and 30-nt were selected. Ribo-seq libraries were then constructed following a protocol described previously [37]. For the 150-μl aliquots lysates, total RNA was isolated with Zymo RNA Clean & Concentrator-25 kit (Zymo Research), RNA-seq libraries were constructed with VAHTS Total RNA-seq (H/M/R) Library Prep Kit (Vazyme, NR603-02) according to the manufacturer’s instructions. All libraries were sequenced with PE150 mode by Illumina Hiseq X10 or NovaSeq 6000.

### Cell cycle analysis

About 1.2X10^5^ cells were seeded in 6-well plates and transfected with siRNA on the following day at around 40% confluence. Cells during exponential growth phase were collected at 24h after siRNA transfection and fixed in 100% methanol at −20°C overnight. Fixed cells were washed twice with PBS, treated at 37°C for 30min with staining buffer containing 25μg/ml propidium iodide (Sigma, P4864), and 50μg/ml RNase A (Invitrogen, EN0531). DNA content was measured by flow cytometry (BD, LSR Fortessa) and analyzed by Modfit software (v4.1).

### Western blotting

Cell lysates were collected at 24h after RP knockdown, denatured at 100℃ for 5min and then separated on 12% PAGE gels for 30min at 70V followed by 1h at 100V. Protein extractions were electroblotted on a polyvinylidene fluoride (PVDF) membrane (Millipore). Membranes were blocked with 5% nonfat dry milk in TBST for 1h and probed with 1:1000 anti-p53 (cst, 2524s), 1:2000 anti-p21 (cst, 2947s) or 1:5000 anti-GAPDH (Proteintech, 60004) primary antibodies. Membranes were then incubated with anti-mouse or anti-rabbit horseradish peroxidase-coupled secondary antibodies at a 1:10000 dilution for 2h. Protein signals were developed with Immobilon Western Chemiluminescent HRP Substrate (Millipore) and imaged with ChemiDoc™ Imaging Systems (Bio rad).

### Cell proliferation and apoptosis analysis

A549 cells transfected with siRNAs were seeded (∼0.6X10^5^ cells/well) and grown in 24-well plates for 0h, 24h, 48h and 72h, respectively. Cell proliferation was assessed by adding 100µl 5mg/ml 3-[4,5-dimethylthiazol-2-yl]-2,5-diphenyltetrazolium bromide (MTT) for 4h at 37°C, followed by aspiration of the medium and addition of 300µl dimethyl sulfoxide. Each reaction was performed in triplicate. Spectrophotometer readings at 490nm were determined with Synergy H1 (Bio Tek). Cell apoptosis was measured by TUNEL staining assay. Dead cells were detected with DNA fragmentation using *In Situ* cell death fluorescein (Roche, 11684795910) following the manufacturer’s instructions.

### Cell senescence detection

A549 cells were seeded in 6-well plates, transfected with siRNA on the following day at around 30% confluence, then trypsinized and counted in triplicate at 72h after transfection. Senescence-associated-*β*-galactosidase (SA-*β*-gal) activity was assayed according to the manufacturer’s instructions (Cst, 9860).

### Preparation of conditioned medium (CM) of *RPL15*-deficient A549 cells

A549 cells were seeded in 12-well plates at a density of 1×10^5^ cells per well with DMEM, transfected with *RPL15*-targeting or control siRNAs using Lipofectamine RNAiMAX reagent according to the manufacturer’s protocol. Medium were replaced with EBM2 containing 0.5% FBS at 6h. Under normoxia, cells were cultured at CO_2_ incubator for 18h. Under hypoxia, cells were cultured in a modular incubator chamber that was flushed with 1% O_2_/5% CO_2_/balance N_2_ at 37°C for 18h; The CMs were harvested, centrifuged at 1000 rpm for 10 min, then stored at −20 °C.

### ELISA assay for VEGF

VEGF concentrations in the CMs from *RPL15*-deficient and control A549 cells under normoxia or hypoxia were detected with Human VEGF Valukine ELISA Kit (Novus, val106).

### Wound-healing assay

HUVECs were seeded in 6-well plates at a density of 2×10^5^ cells per well, incubated until an even monolayer reached around 90% confluence. HUVEC monolayers were scraped with a 200-μL pipet tip. Cells were washed to remove detached cells with PBS before incubation with CMs. The healing wounds were photographed at 0h and 8h, respectively.

### Tube formation assay

A 48-well plate was paved with growth factor reduced matrigel (Corning, 354230) and maintained at 37°C for 30min to make the matrix solution to gel. HUVECs were seeded on the gel and cultured in CM for 48h at 37 °C with 5% CO_2_. To assess the tube formation and disassembly, cells were photographed at 6h, 12h, 18h, 24h, 30h and 48h, respectively. Tube numbers and lengths were evaluated by Image J software [38]. Each experiment was repeated for six times.

### shRNA plasmids and RNAi interference of RPs in mouse retinas

*In vivo* conditional knockdown of *RPL11* or *RPL29* were performed in retinas of neonatal mice. For knockdown experiments, selected small hairpin sequences were inserted into the shRNA interference vector pBS/U6 containing the human U6 promoter [39]. Primer sequences for constructing pBS/U6-shRNA plasmids are shown in Table S1. Knockdown efficiency was tested by qRT-PCR in neuro-2a cells. The constructs with high knockdown efficiency were used in the following experiments. The targeting sequence for mouse *RPL11 (mRPL11)* is: 5’-CCGCAAGCTCTGCCTCAATAT-3’ (*mRPL11*-sh), for *mRPL29* is: 5’-GCCAAGAAGCACAACAAGAAA-3’ (mRPL29-sh) and for non-targeting shRNA control (shNT) is: 5’-GCGCGATAGCGCTAATAATTT-3’. To perform retinal knockdown, pBS/U6 constructs and pCIG vector (as a GFP reporter) were mixed at a ratio of 2:1 (μg/μl). 1 μl mixture was injected into the subretinal space of P0 CD1 mice with a Microliter Syringe (Hamilton). Immediately following injection, electric pulses (100 V; five 50-ms pulses with 950-ms intervals) were applied with the ‘‘+’’ electrode of tweezer-type electrodes (BTX) positioned on the injected eye. Transfected retinas were collected at P12 for analysis when the great majority of retinal cells were determined and developed into mature cell types.

### Polysome profiling

Polysome profiling was conducted following a previous study [40]. In detail, one 10cm dish of A549 cells were treated with 100μg/ml CHX in DMEM at 37°C for 5min prior to harvest and then washed twice with cold PBS containing 100 μg/ml CHX. Samples were lysed with 800μl lysis buffer containing 10mM Tris-HCl pH 7.4, 5mM MgCl_2_, 100mM KCl, 1% Triton X-100, 2mM DTT, 100μg/ml CHX, Complete Protease Inhibitor EDTA-free (Roche, 4693132001) and 20 U/mL SUPERase In RNase inhibitor, scraped and transferred to eppendorf tubes, then kept on ice for 10min. Lysates were centrifuged at 10000xg for 10min at 4°C. RNA concentrations were measured with Nanodrop UV spectrophotometer (Thermo Fisher Scientific). Equal amounts of lysates were layered onto a liner sucrose gradient (10%∼50%, w/v) with the gradient buffer containing 20mM HEPES-KOH pH 7.4, 5mM MgCl_2_, 100mM KCl, 2mM DTT, 100μg/ml CHX, and 20 U/mL SUPERase In RNase inhibitor, centrifuged in a SW41 Ti rotor (Beckman) for 2h at 36000rpm at 4°C. Samples were fractioned and analyzed by gradient station (BioCamp).

### Construction and screening of lentiviral sgRNA library of human RPs

sgRNA sequences (Table S7) for human RP genes were retrieved from Brunello library [41]. Four sgRNAs for each RP and 80 non-targeting sgRNAs were designed and synthesized with a CustmoArray 12K array chip (CustmoArray, Inc.). Plasmid library was constructed following a previous study [41] with minor modification. sgRNA libraries were amplified as sub-pools in a nested PCR. For the first round of PCR, all sgRNAs were amplified using Phusion Flash High-Fidelity PCR Master Mix (NEB, M0531L) with the primers: sense: 5’-ACGCTCAGTTCATATCATCACG-3’, antisense: 5’-ATCGCAGCATCTACATCCGATGT-3’. The second round of PCR was performed to incorporate overhangs compatible into lentiCRISPRv2 vector using the primers: sense: 5’-TTTCTTGGCTTTATATATCTTGTGGAAAGGACGAAACACCG-3’, antisense: 5’-GACTAGCCTTATTTTAACTTGCTATTTCTAGCTCTAAAAC-3’.

Meanwhile, lentiCRISPRv2 plasmid was digested using *BsmB*I and purified. The digested plasmids and sgRNAs were ligated using Gibson Assembly Master Mix and transformed into competent DH5α cells. The number of clones for each sgRNA was about 400 in average.

The lentivirus was produced by co-transfection of lentiCRISPRv2-sgRNA-RPs with the pVSVg and psPAX2 into 293T cells using Lipofectamine 3000 reagent (Invitrogen, L3000). Medium were changed at 6h after transfection. The virus-containing supernatant was collected 48h after transfection and filtered by a 0.45 μm filter. A549 cells were infected with sgRNA library lentiviruses at an MOI of < 0.3. Genomic DNA extraction were conducted in 2 batches: the first batch of cells (5×10^6^) were collected 2 days after infection using the TIANamp genomic DNA kit (Tiangen, DP304), and the second batch of cells treated with puromycin were collected at 7d. For each library, the lentiviral integrated sgRNA-coding regions were PCR-amplified using sense: 5’-AATGGACTATCATATGCTTACCGTAACTTGAAAGTATTTCG-3’, antisense: 5’-TCTACTATTCTTTCCCCTGCACTGTtgtgggcgatgtgcgctctg-3’ by TransTaq HiFi DNA polymerase at first, and the sequencing libraries were amplified with Titanium Taq (Clontech, 639209) with the following barcoded primers for control sample: sense: 5’-AATGATACGGCGACCACCGAGATCTACACTCTTTCCCTACACGACGCTCTTCCGATCTtAAGTAGAGtcttgtggaaaggacgaaacaccg-3’, antisense: 5’-AAGCAGAAGACGGCATACG AGATAAGTAGAGGTGACTGGAGTTCAGACGTGTGCTCTTCCGATCTtTCTACTATTCTTTCCCCTGCACTGT-3’, and with the following barcode primers for experiment samples: Sense: 5’-AATGATACGGCGACCACCGAGATCTACACTCTTTCCCTACACGACGCTCTTCCGATCTatACACGATCtcttgtggaaaggacgaaacaccg-3’, antisense: 5’-CAAGCAGAAGACGGCATACGAGAT ACACGATCGTGACTGGAGTTCAGACGTGTGCTCTTCCGATCTatTCTACTATTCTTTCCCCTGCACTGT-3’. The products were further purified with Gel Extraction kit (Tiangen, DP209) and prepared for sequencing by Illumina Novaseq 6000.

### Quantification and assessment of sgRNA effects in CRISPR experiment

Sequenced reads containing specific sgRNA sequences were grouped and counted. The sum of all the counted reads for each library was taken as the library size. Normalized sgRNA expression level was calculated by dividing sgRNA counts by corresponding library size. Minimal ratio of sgRNA expression level at day7 vs day2 was defined as the repression effect on cell growth of corresponding RP upon knockdown. For quality control, normalized sgRNA expression at day7 and day2 were compared. Theoretically, non-targeting sgRNAs would increase at day7 while RP-targeting sgRNAs would decrease at day7. The repression effect for each RP in our assays correlated (PCC R = 0.46, *P* = 3.8e-05) with that in the Cancer Dependency Map Project (DepMap) database (https://depmap.org/portal/) [42].

### Sequencing data processing and quality control for RNA-seq and Ribo-seq

Base callings were demultiplexed and converted to fastq files with Bcl2fastq (v2.20.0.422) (https://support.illumina.com/downloads/bcl2fastq-conversion-software-v2-20.html). Adapters and low-quality reads ending with quality scores < 20 were trimmed by Cutadapt (v1.8.1) (http://code.google.com/p/cutadapt/). Trimmed reads with length < 20nt were discarded. Reads aligned to human rRNA and tRNA sequences extracted from the UCSC database (http://genome.ucsc.edu/) [43] were further excluded. For Ribo-seq libraries, trimmed reads with length < 26nt or > 34nt were excluded. Remaining reads were aligned to the human genome (Ensembl GRCh38.88) [44] with STAR (v2.6.1-d) (https://github.com/alexdobin/STAR) [45].

For quality control, we assessed each library from multiple aspects. (a) Total and unique alignments were summarized using the resulting files by STAR. (b) Gene body coverage was estimated to test whether sequence bias occurred in 3’ or 5’ end of transcripts by “*geneBody_coverage.py*” from RSeQC [46]. (c) Relative percentages of alignments to gene features (CDS, 5’ UTR and 3’ UTR of annotated transcript according to GTF file) were estimated by “*read_distribution.py*” from RSeQC. For gene body coverage and read distribution analyses, we included transcripts with combined exon length >1000nt and at least two exons. (d) For Ribo-seq libraries, frame distributions and ribosomal footprints around the start or stop codons were analyzed by “*ribotish quality*” with default parameters, embedded in Ribo-Tish (v0.2.2) [47].

### Gene expression quantification, normalization, and differential analyses

To allow for proper comparison and integration of mRNA-seq and Ribo-seq data, all RNA-seq quantifications were derived from the first reads mapped to the defined genomic features in Ensembl GTF file as previous study [48]. Unique alignments to exons in RNA-seq and that to CDS in Ribo-seq were counted with featureCounts (v1.6.0) [49] for quantification of gene expression. Raw gene counts were converted to RPKM (short for Reads Per Kilobase per Million mapped reads) values following the formula:

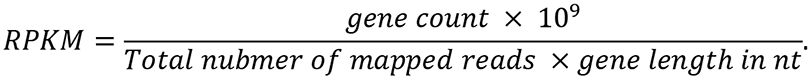

For RNA-seq and Ribo-seq datasets, we included the genes with mean RPKM > 1 across all libraries in further analysis (*n* = 13,294 for RNA-seq; *n* = 11,694 for Ribo-seq). We set up replicates for 16 RPs and the replicates showed highly reproducible, we thus combined the replicates using the mean RPKM values to define gene expression. Our sequencing datasets included hundreds of libraries, prepared in several batches. Batch effects in RNA-seq and Ribo-seq were evaluated and eliminated by ComBat [50] in ‘sva’ package [51]. Differential expression analysis was performed by comparing normalized gene expression in RP-deficient cells with that in control cells from the same batch. For both RNA-seq and Ribo-seq data, we defined a differential gene when its fold change ≤ 0.5 or ≥ 2.

### Reproducibility analysis

To test the reproducibility of our datasets, we firstly compared genome-wide gene expression levels between replicates using Pearson correlation analysis. We also compared gene expression in our control cells to public datasets. In addition, we compared the functional associations in our analysis with the previous studies by extensive literature search.

### Clustering analysis of differential genes

Differential genes for each RP were combined and clustered to detect common or distinct signals for downstream gene expression changes between RPs. In detail, indicator matrices containing values −1, 0 and 1 for RNA-seq and Ribo-seq were used, where each row represents a gene, and each column represents a specific RP. For indicator matrix, 1 referred to the upregulation, −1 referred to downregulation, and 0 referred to the unchanged genes upon knockdown of indicated RP. For RP clustering, hclust was performed using the correlation matrix between RPs as input. For gene clustering, kmeans was performed. Cluster visualization was done using “Heatmap” from ComplexHeatmap package [52].

### Functional enrichment analyses

For each RP, GO enrichment [53, 54] and KEGG pathway [55, 56] were conducted with gprofiler2, an interface to the g:Profiler toolset (https://biit.cs.ut.ee/gprofiler/gost) [57]. The up-regulated or down-regulated genes were used as input for each RP. To remove the redundancy between GO terms, hclust algorithm was performed according to the similarity measurement by ‘GOSemSim’ [58, 59] and GO classes were determined by “silhouette” method.

### Translational efficiency estimation and differential analysis

Translational efficiency (TE) estimations were calculated by taking the ratio of gene expression level from Ribo-seq over that from RNA-seq data. In detail, for RNA-seq and Ribo-seq libraries, we only counted the unique alignments that fully contained within the CDS regions thus avoiding counting reads that overlap multiple features. Raw gene counts were then converted to RPKM values. Batch effects were also assessed and excluded. Normalized gene expression matrices for RNA-seq and Ribo-seq were then used to calculate TE. Differential analysis was performed by comparing the TE in RP-deficient cells with that in the control cells from the same batch.

### Dissecting transcriptional and translational regulation

RPFs in Ribo-seq data reflect the combined outcome of transcriptional and translational control in gene expression. A gene’s regulation mode was defined by intersecting the changes in RNA-seq (DEGs), Ribo-seq (DTGs) and TE level (DTE). In total, four classes of regulation mode were defined including forwarded, exclusive, intensified and buffered regulation, where forwarded genes indicated that Ribo-seq changes are explained by RNA-seq changes; Exclusive genes indicated that TE changes occur exclusively without underlying mRNA changes; Buffered and Intensified genes indicated that both the TE and the mRNA are changing.

### Other datasets

Quantification of P53 protein levels and the impact on nucleolar disruption which was estimated by quantification of nuclear stress (iNO index) after RP deficiency were extracted from Emilien Nicolas *et al*’s study [11]. In brief, they used specific siRNAs to deplete RP expression one-by-one in FIB364 or HCT116 cells. A549 RNA-seq data from CCLE was extracted by “depmap” package.

### Defining cell type-specific genes in A549 cell

To identify the cell type-specific expressed genes in human cancer cell lines, RNA-seq-based gene expression quantification for 1165 cancer cell lines were extracted using “depmap” package, an agent for the Cancer Cell Line Encyclopedia (CCLE) database [60]. Aggregated log2-scaled TPM values were used. We defined the genes with TPM values > Mean+2SD as cell type-specific genes for a certain cell type. We totally identified 478 A549-specific expressed mRNAs, out of which 199 were protein-coding genes. We focused on the protein-coding genes in our further analyses. Fisher’s exact tests were used to check whether the differential genes for each RP were enriched by A549-specific expressed protein-coding genes.

### Extraction and characteristics of transcript features

Nucleotide sequences for genome-wide or individual transcript features, such as 5’ UTR, were extracted from the Ensembl database [44]. Transcripts with short 5’ UTRs (< 20 nt) or long 5’ UTR (> 500nt) [31] were excluded in further analysis. To quantify mRNA structure complexity, the minimum free energy (MFE) was predicted for selected transcripts with RNAfold (https://www.tbi.univie.ac.at/RNA/RNAfold.1.html) [61]. The GC content and length of mRNAs of interest were estimated with ‘seqinR’ package (http://r-forge.r-project.org/projects/seqinr/) [62]. Kozak sequence was defined using the sequence 10nt upstream of start codon for each transcript. Kozak sequence similarity analysis and visualization were performed with motifStack [63] and ggseqlogo [64] package respectively.

### Quantification and statistical analysis

Statistical information including n, mean, and statistical significance values are indicated in the text or the figure legends. Results were statistically analyzed using Student’s t test or an analysis of variance (ANOVA) test with multiple comparisons where appropriate using Prism 8.0 (GraphPad Software, Inc). A *P*-value of < 0.05 was considered to be statistically significant.

## Data and software availability

All sequencing data were deposited to the Gene Expression Omnibus (GEO) with an accession number XXX which will be publicly available upon the acceptance of the manuscript.

## Acknowledgement

We would like to thank Dr. Feng Zhang for providing helpful suggestions. We would also like to acknowledge the support from the Center for Precision Medicine, Sun Yat-sen University.

## Author Contributions

Z.X. conceived and supervised the study. N.T. and JQ.Y. prepared sequencing libraries and performed high-throughput sequencing experiments. YZ.L. performed sequencing data processing, quality assessment and analysis. Y.W. and HW.W. helped sequencing data analysis. R.J., F.Z., N.T. and YZ.L. designed experiments on *RPL15*. N.T. and JQ.Y. performed experiments on *RPL15*. R.J. and MQ.X. supervised experiments on *RPL15* and *RPL11*. YZ.L., N.T. and ST.L. designed mouse retina experiments. ST.L. and YN.G. performed mouse retina experiments. CC.C. helped quality control analysis of sequencing datasets. YZ.L., N.T., JQ.Y., ST.L. and R.J. prepared the materials for the manuscript. YZ.L. and Z.X. wrote the manuscript. All authors read and approved the manuscript.

## Supplementary Figures and Tables

**Figure S1.**
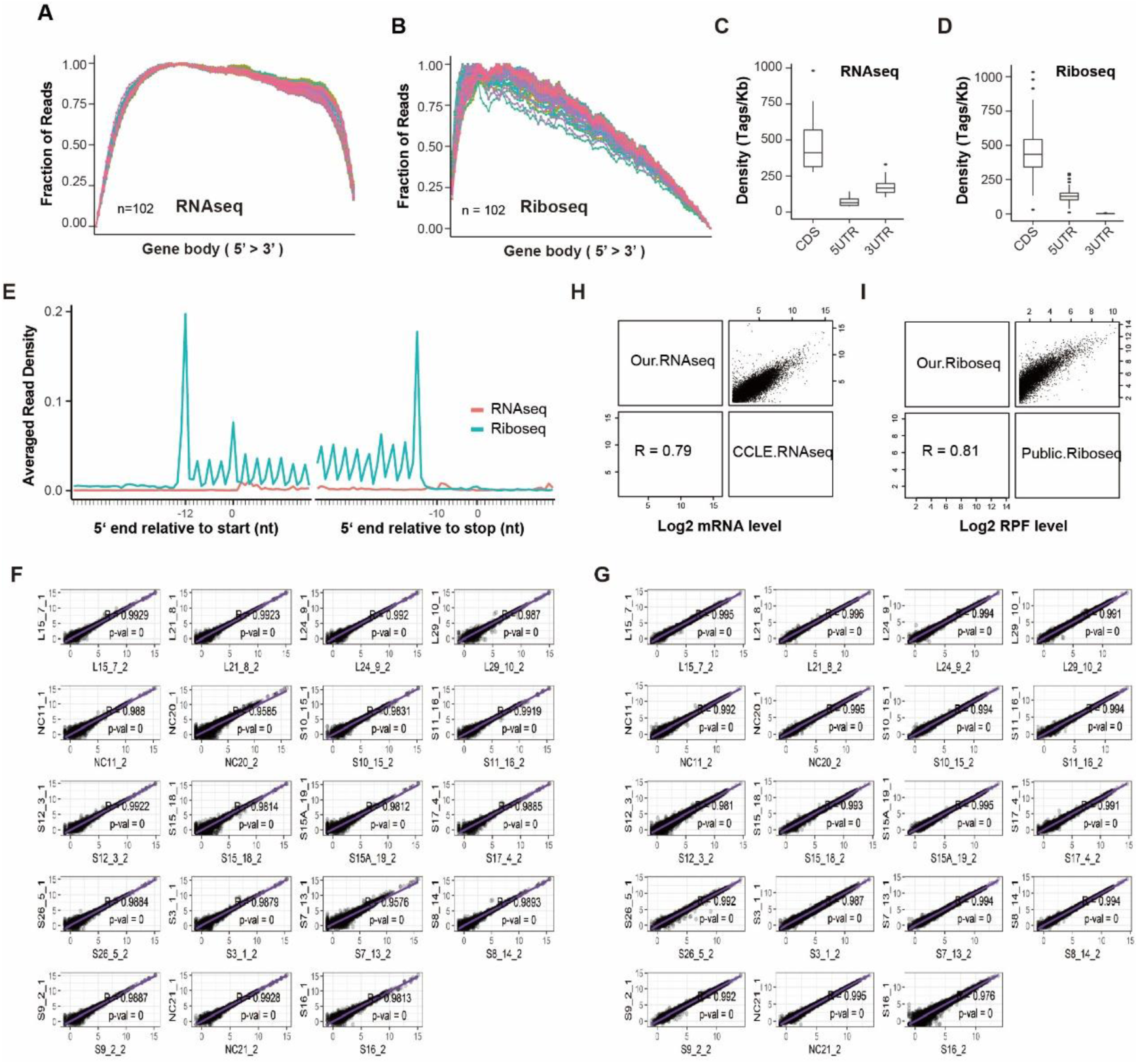
Overview of the transcriptome and translatome datasets, related to Figure 1. (A, B) Gene body coverage (5’-3’) for each library in RNA-seq (A) or Ribo-seq (B). Each line indicates one sample. (C, D) Relative density of short reads mapped to CDS, 5’ UTR and 3’ UTR in all libraries from RNA-seq (C) and Ribo-seq (D). (E) Representative density plots showing comparison of read density along each nucleotide around the start codons and stop codons in RNA-seq and Ribo-seq of one selected RP. (F, G) Pearson correlation between libraries from replicates in RNA-seq (F) or Ribo-seq (G). Log2 RPKM values were used. (H, I) The correlation analysis between our control A549 cells and public A549 datasets by RNA-seq (H) or Ribo-seq (I). Log2 RPKM values were used in correlation analysis.

**Figure S2.**
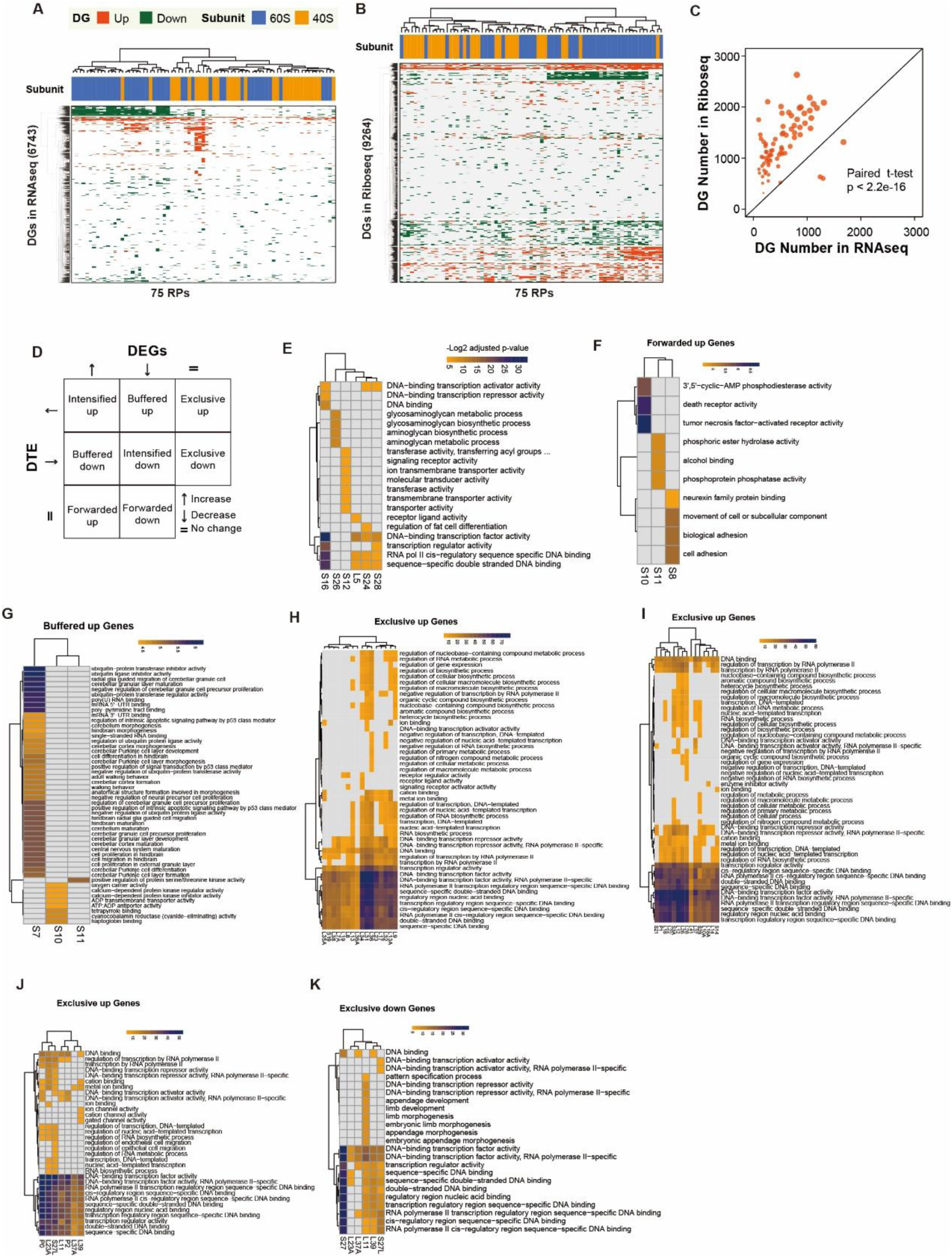
Diversity of gene expression changes upon RP knockdown, related to Figure 2. (A, B) The clustering of RPs according to common DEGs (A) or common DTGs (B). 6743 DEGs and 9264 DTGs were included. For clustering analysis, numeric matrix containing −1, 0, and 1 for RNA-seq and Ribo-seq are used, where −1 indicated down-DGs, 1 indicated up-DGs and 0 indicated non-changed genes for a RP. Hierarchical clustering analyses were performed for both rows (genes) and columns (RPs). (C) The comparison of numbers of DEGs and DTGs for each RP. Paired T-test was performed. (D) Intersecting DEGs and DTE genes showing several categories of gene expression regulation mode for each RP. Forwarded genes, where the Ribo-seq changes are explained by the RNA-seq changes; Exclusive, where changes occur exclusively in TE without underlying mRNA changes; Buffered and Intensified, where both the TE and the mRNA were changing. (E-K) The enriched GO terms by differential genes under primary regulation mode for indicated RPs as shown in Figure 2E.

**Figure S3.**
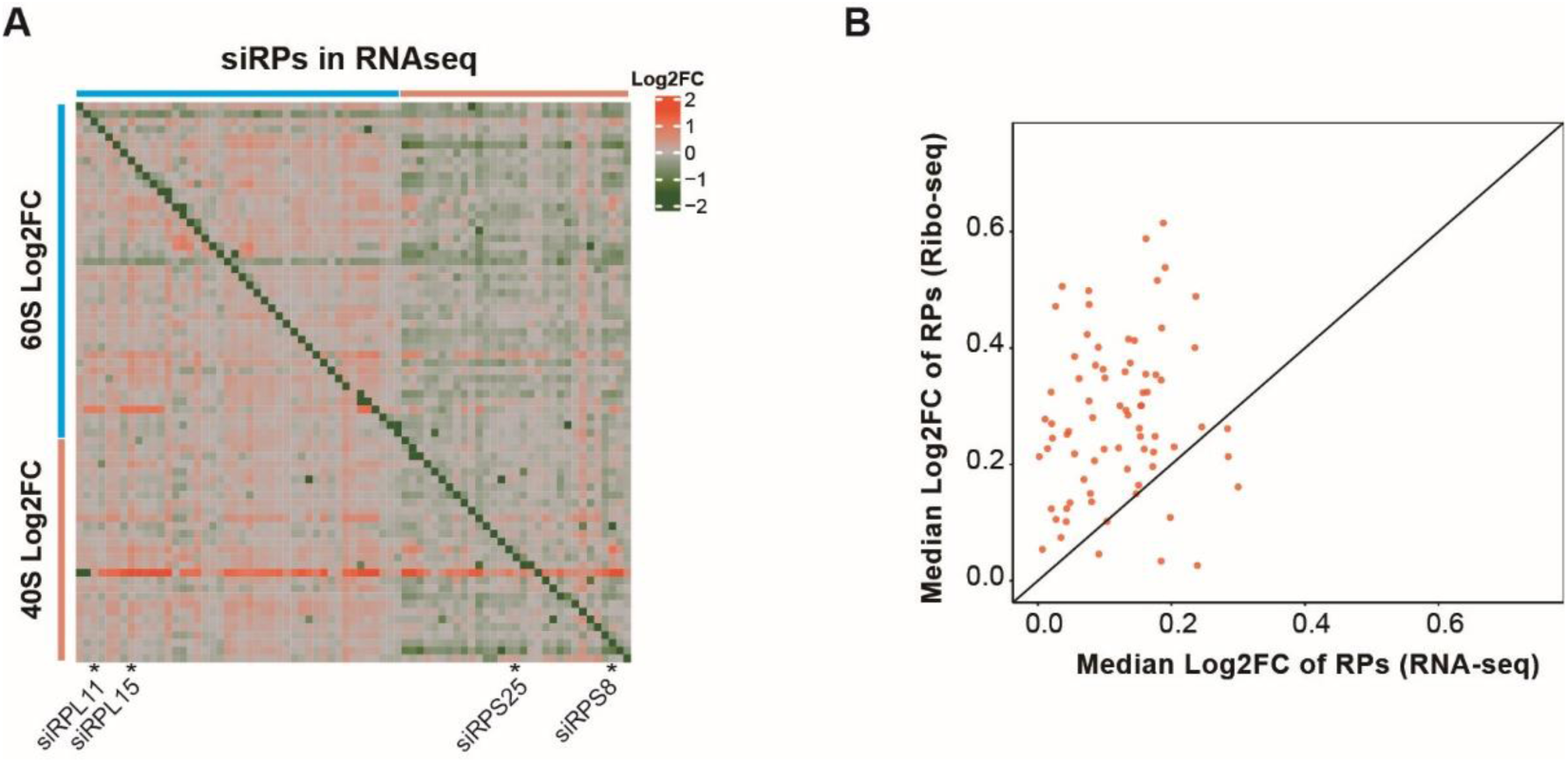
co-regulation of 60S and 40S RPs after RP deficiency, related to Figure 3. (A) The gene expression changes of 60S and 40S RPs in RNA-seq after knockdown of RPs from 60S or 40S. (B) The comparison of gene expression changes of other RPs between RNA-seq and Ribo-seq. For each targeted RP, the median Log2FC values of other RPs were used.

**Figure S4.**
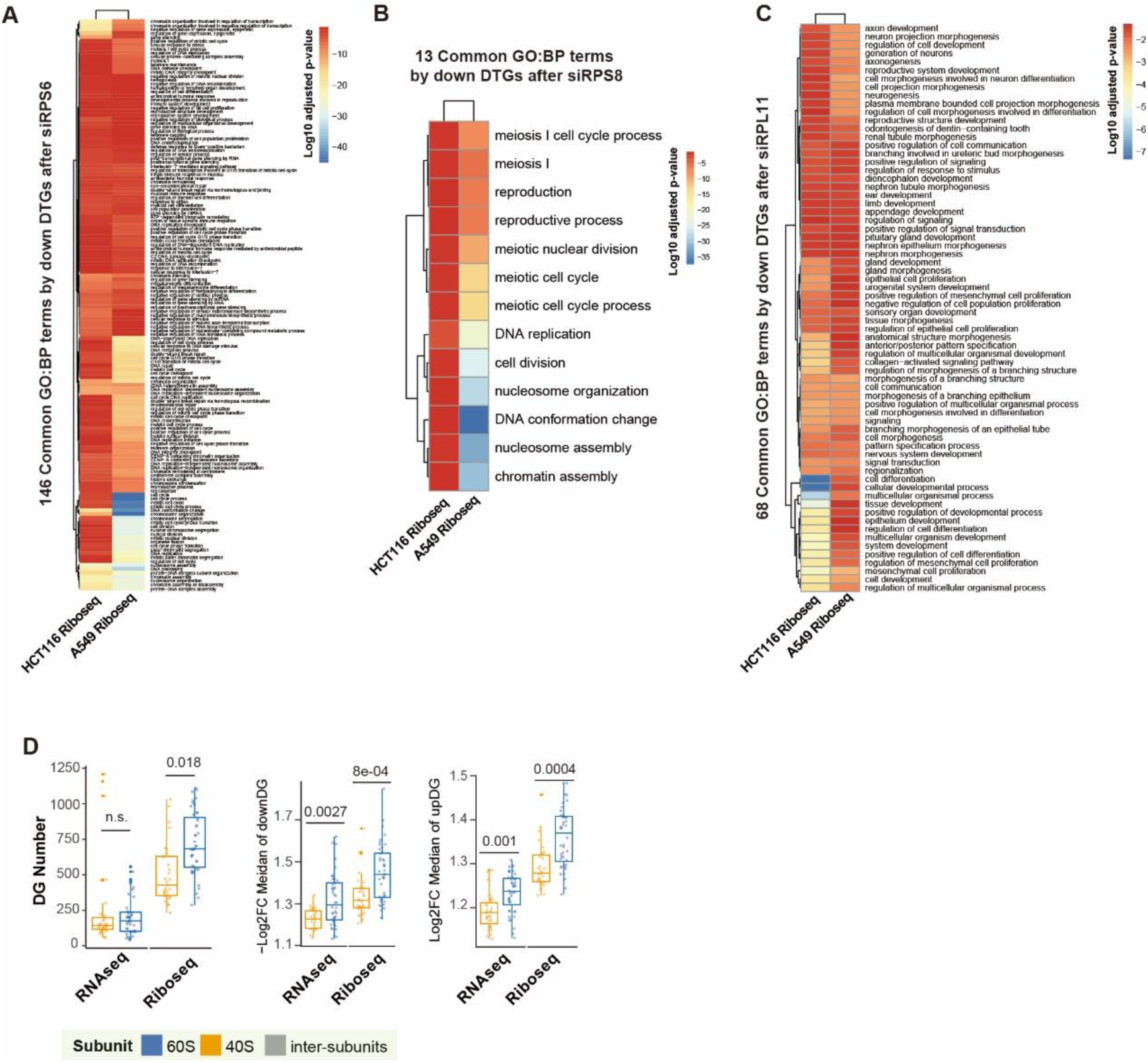
Functional characterization of RPs, Related to Figure 4. (A-C) The significance of overlapped enriched BP terms by downregulated DTGs after knockdown of *RPS6* (A), *RPS8* (B) or *RPL11* (C) in HCT116 cells and A549 cells. (D) The comparisons of DG numbers (left panel), median fold change of down-DGs (middle panel) or up-DGs (right panel) after knockdown of 40S or 60S RPs in RNA-seq and Ribo-seq. T-tests were performed.

**Figure S5.**
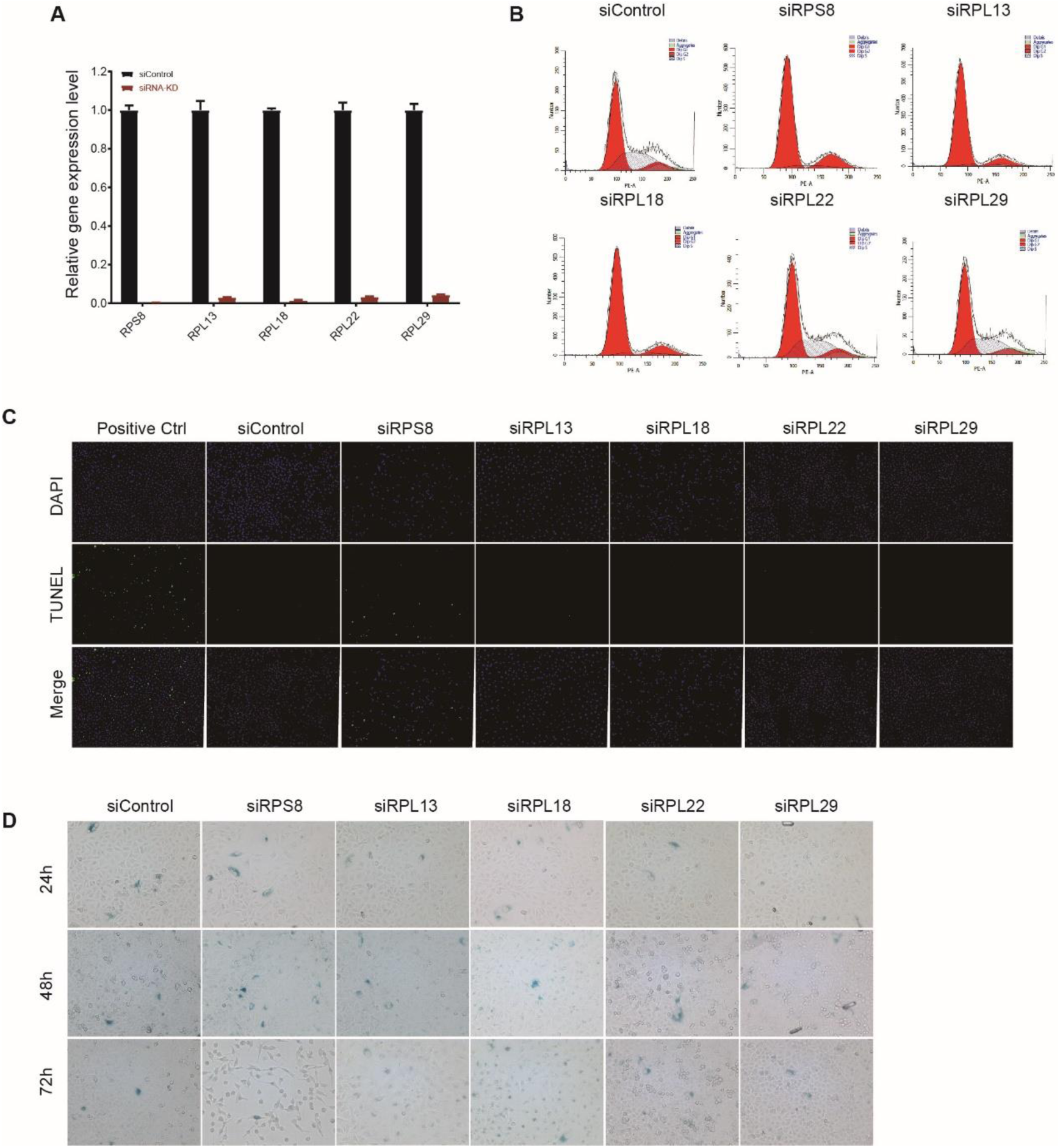
RP deficiency promoted different cell fates after cell cycle arrest, related to Figure 5. (A) Bar plots showing high efficiency of knockdown by specific siRNAs for *RPS8*, *RPL13*, *RPL18*, *RPL22*, or *RPL29*. mRNA levels in control and treated cells were quantified by qPCR. For each RP, mRNA levels relative to control samples are shown. (B) Representative of propidium iodide (PI) staining and flow cytometry for cell-cycle analysis in A549 cells treated by siRNAs without targets (control) or targeting indicated RPs. (C) Representative of TUNEL staining for testing apoptosis *in situ* in A549 cells after knockdown of *RPS8*, *RPL13*, *RPL18*, or *RPL22*. (D) Representative of β-Galactosidase (β-Gal) staining assays for testing senescence in A549 cells after knockdown of *RPS8*, *RPL13*, *RPL18*, *RPL22*, or *RPL29*.

**Figure S6.**
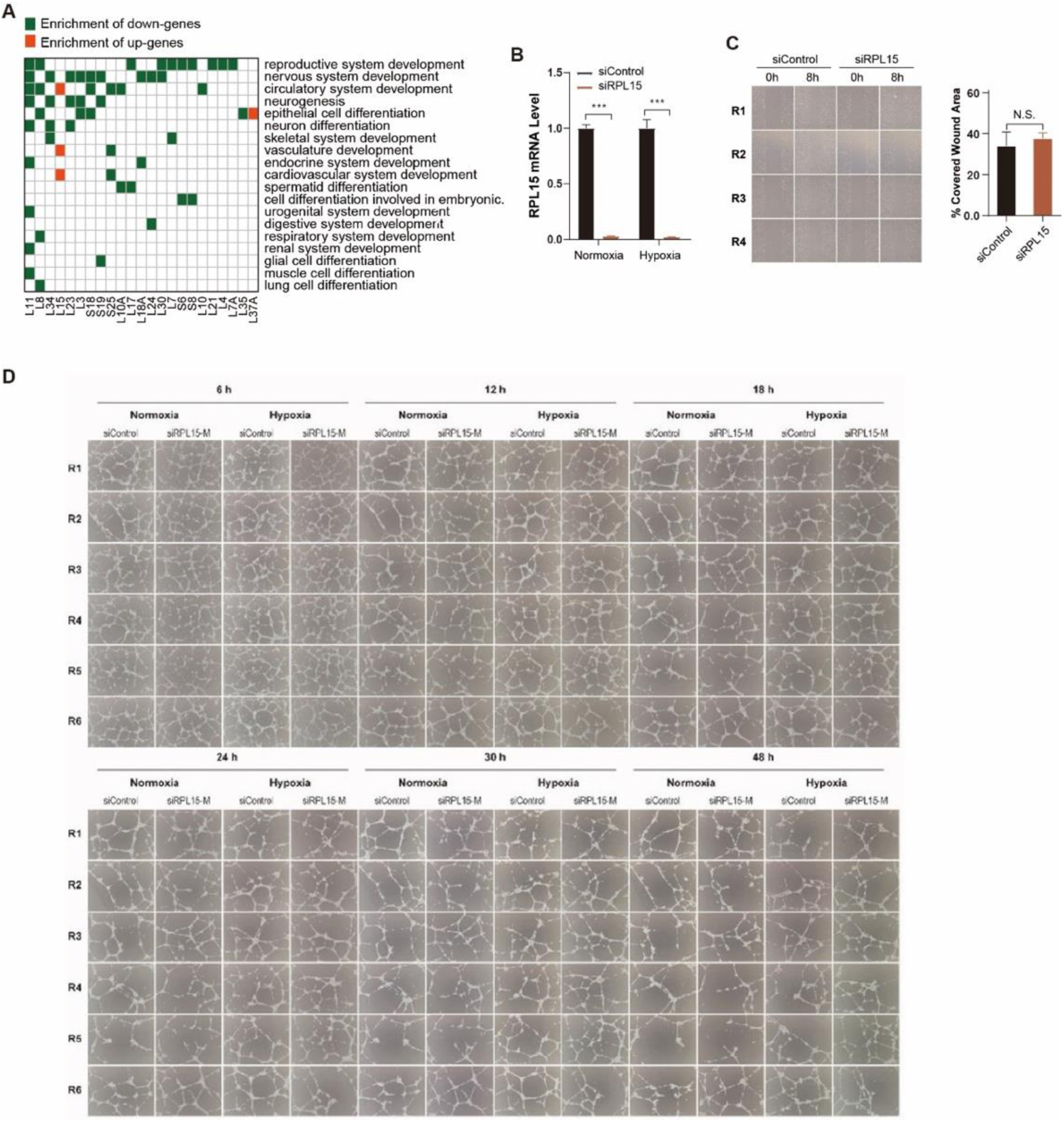
RP deficiency impacts tissue development, Related to Figure 6. (A) Heatmap showing the enrichment of system development or cell differentiation by up- or down-DGs after knockdown of RPs in Ribo-seq. (B) Bar plots showing high knockdown efficiency of *RPL15* siRNAs under normoxia and hypoxia environments. *RPL15* mRNA levels were quantified by qPCR. (C) Analysis of the effects on cell migration of the conditioned media (CM) of *RPL15*-deficient A549 cells. (Left panel) Images of HUVEC cells treated by the CM from *RPL15*-deficient A549 cells and the control cells 0hr and 8hr after a scratch was introduced in the monolayer with a pipette tip. (Right panel) The percentage of open wound area for each of four replicates at 8hr over that at 0hr was estimated and compared between the CMs from *RPL15*-deficient cells and the control cells. T-test was used. (D) Images of HUVEC cells in tube formation assays treated by the CM from *RPL15*-deficient A549 cells and the control cells under normoxia and hypoxia environments. HUVEC cells at 6hr, 12hr, 18hr, 24hr, 30hr and 48hr during the assays were photographed. Six replicates were introduced.

**Figure S7.**
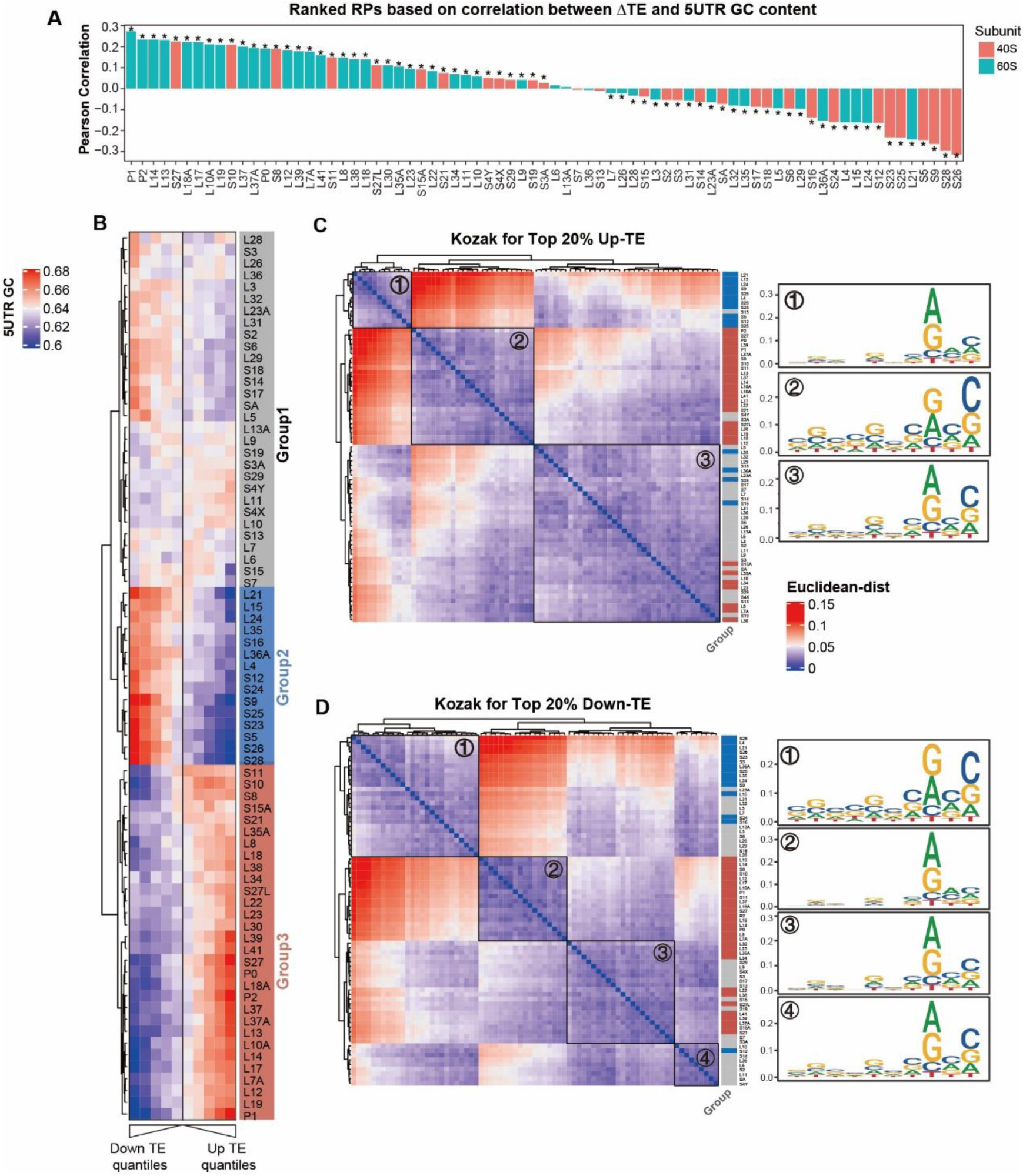
Correlating transcript features and TE changes after RP deficiency. (A) The ranked RPs according to the correlation between global TE changes and 5’ UTR GC content. (B) Heatmap showing the mapping of 5’ UTR GC contents in genes within different quantiles of global TE changes. RPs were clustered by hclust method. (C, D) The clustering of RPs according to the similarity of Kozak sequence of top 20% genes with upregulated TE (C) or downregulated TE (D). Merged kozak context for each subcluster is shown on the right panels.

**Figure S8.**
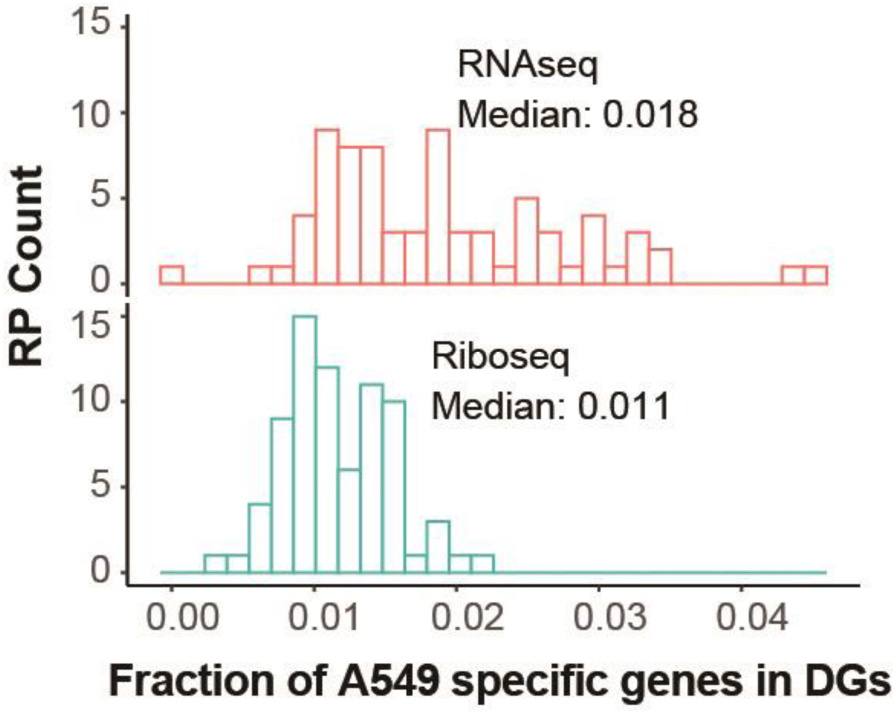
Percentages of A549-specific expressed genes in DEGs or DTGs. Histograms showing the percentages of A549-specific expressed genes in the DGs identified in our RNA-seq and Ribo-seq datasets. A549-specific genes were estimated following the description in “Material and Methods”.

**Table S1. Sense and antisense sequences for siRNAs targeting specific RPs.**

**Table S2. Knockdown efficiency of RPs**. The mRNA changes of RPs relative to the control cells treated by non-targeting siRNAs were estimated and averaged in replicates. The mRNA levels were quantified 24h post treatment with siRNAs.

**Table S3. Mapping statistics for each library of RNA-Seq and Ribo-Seq.**

**Table S4. Functional enrichment by differential genes in RNA-seq and Ribo-seq**. Gene-Set Analyses (GSA) of up-regulated or down-regulated genes in RNA-seq or Ribo-seq for each RP was performed using functional annotation gene sets obtained from gprofiler database.

**Table S5. Recovery of known functional associations of RPs**. Functional associations for each RP were retrieved by literature search, and the overlapped entries between the enriched terms by our sequencing data and the ones reported by at least one studies were included.

**Table S6. qRT-PCR primers for human RP.**

**Table S7. The sgRNA sequences in CRISPR-Cas9 experiments**. Four gRNA sequences for each RP were designed.

